# Single-cell analysis reveals a universal pericyte signature associated with poor clinical outcome and immune T cell dysfunction in thyroid cancer and other cancers

**DOI:** 10.64898/2025.12.07.692700

**Authors:** Aarav Bhasin, Yubin Na, Jack Lawler, Carmelo Nucera

## Abstract

**Background:** Cancer is a devastating disease with rising incidence rates and generally poor outcomes. Single-cell genomic technologies enabled the profiling of thousands of individual cells from tumors to understand their role in cancer progression. In this study, we are characterizing pericytes from tumor microenvironment single-cell data of human cancers to assess associations with patient prognosis and achieve mechanistic insights.

**Design:** we have characterized pericytes from tumor microenvironment (TME) single-cell datasets of different cancers to assess associations with patient prognosis and achieve mechanistic insights.

**Methods:** For comprehensive pericyte characterization, 23 publicly available single-cell RNA sequencing (scRNA-seq) datasets corresponding to 15 cancers and 1 benign tumor were downloaded from the Gene Expression Omnibus (GEO) database and Tisch repositories. The datasets were processed using a uniform workflow, including quality control, normalization, variable gene selection, clustering, and cellular annotation (based on automated and manual markers). The annotation of the pericytes was performed based on the score calculated using our previously validated pericyte gene signature. The differential gene expression analysis between pericytes and fibroblasts in each dataset was performed to identify pericyte signature. Comparative analysis of the signatures was performed to identify a universal pericyte signature that was evaluated for pericyte specificity using peripheral blood mononuclear cells (PBMC) data. The universal pericytes signature genes were evaluated for cancer outcome associations in The Cancer Genome Atlas (TCGA) datasets using the Survival Genie Platform. Furthermore, pathway and gene-network analyses were performed to understand the biological significance of the pan-cancer universal pericyte signature.

**Results:** We conducted an analysis of 23 single cell RNA-sequencing datasets from 14 cancers: breast, pancreatic, bladder, ovarian, non-small cell lung, cervical, prostate, basal cell, colon, head and neck, papillary and anaplastic thyroid cancers, as well as melanoma and lymphomas; and a benign tumor (neurofibroma). The pericytes gene set enabled the identification of a distinct pericyte cluster comprising more than 50 cells in the majority of datasets. Differential gene expression analysis between pericytes and fibroblasts identified heterogenous pericyte signatures for different cancers. Initial cell–cell communication analysis in anaplastic thyroid cancer indicated that pericytes are highly communicative cells, ranking among the top ligand-secreting populations, and actively engaging with anaplastic thyroid cancer cells and T cells. A comparative analysis of pericyte signatures generated from different cancers yielded a core signature of 100 genes that were consistently overexpressed in more than 60% of the cancer datasets. Notably, 78% of these genes were not expressed in PBMCs, supporting their pericyte specificity. Survival analysis using TCGA datasets identified a 19 genes pan-cancer pericyte signature associated with poor prognosis (HR>1 in at least 30% datasets). This pan-cancer pericyte signature included genes such as TPM2 and EHD2, whose overexpression was significantly associated with poor overall survival across multiple aggressive cancers, including bladder, head and neck, lung, pancreatic, skin, and thyroid cancers. Interestingly, many of these genes showed a strong positive correlation (e.g. PDGFRB) with T cell exhaustion-related genes in the TCGA pan-cancer cohort (n=10,293 samples), suggesting a potential mechanistic link between pericytes and T cell exhaustion. Importantly, we have also identified a unique tyrosine kinase (TK) receptors gene signature in pericytes compared to fibroblasts or other cell types by heatmap analysis across the different cancers.

**Conclusions:** This large-scale scRNA-seq analysis of multiple datasets defines a pan-cancer pericyte signature and sheds light on pericyte activity, particularly their communication with T cells, suggesting a role in T cell exhaustion, a key factor in the failure of many cancer therapies.

This study underscores the pivotal role of pericyte population enhancement as a critical regulator of the TME, facilitating immune evasion and driving disease progression. TK expression in pericytes may serve as a predictive biomarker for TKI responsiveness, offering a cellular target whose expression pattern may guide therapeutic decisions and improve patient stratification in TKI-based treatment regimens.

**Translational significance:** These findings provide clinically actionable insights by identifying pericyte-derived molecular targets that may refine patient risk stratification and support personalized treatment approaches. The pericyte-associated markers uncovered here show strong prognostic value across multiple cancer types, highlighting their potential to improve outcome prediction. Moreover, the specificity of these targets positions them as promising candidates for therapeutic intervention and for incorporation into future biomarker-driven clinical trials.

## Introduction

In the recent two decades, a decline in cancer-related mortality was observed, largely due to screening programs and improvements in treatment protocols ^1^. The introduction of targeted and immunotherapies therapies played a key role in this decline, particularly in hematologic cancers and melanoma ^2,3^. However, resistance mechanisms are frequently observed, and cancer still caused deaths of over 600,000 people in 2024 ^1^. Thyroid cancer is the most common malignancy among endocrine organs and ranks as the 7^th^ most common cancer worldwide, according to data coming from the World Health Organization ^4,5^. Its incidence has been steadily increasing over time and is projected to become the fourth most common cancer by 2030 ^6^, with an estimated 2.4-fold increase since the early 2000 ^7^. While this rise has been largely attributed to improved diagnostic capabilities, including the detection of small or moderately small tumors, there is also a growing incidence of aggressive cases^6^. Radioiodine resistant cases as well as poorly differentiated thyroid cancer (PDTC) and anaplastic thyroid cancer (ATC) cases have a poor survival rate ^8–10^. Tumor microenvironment (TME) plays a pivotal role in disease progression and therapeutic resistance, and its modulation has emerged as a promising strategy for targeted therapies ^11–13^.

We have previously observed that pericytes are key components of the thyroid TME ^13–15^. Pericytes are also abundant in other tumors such as ovarian and breast cancer, melanoma and lymphoma ^16,17^. Together with endothelial cells, pericytes sustain vasculature structure and regulate angiogenesis ^18^. They communicate with endothelial cells by paracrine and physical signals promoting vascular formation and remodeling^19^. Furthermore, immunosuppressive effects have been recently described in glioblastoma model, where pericytes secrete anti-inflammatory cytokines and immunosuppressive molecules after cancer cell signals ^20,21^. We have previously showed that pericytes modulate the therapeutic response in papillary (PTC) and ATC derived cells *in vitro* and *in vivo* ^13–15^. Pericytes have been associated with increased melanoma cell growth ^22^. Historically, pericytes absence was associated with worse prognosis in colorectal cancer patients by immunohistochemistry experiments ^23^. However, new evidence driven by single-cells RNA sequencing (scRNA-seq) showed that pericytes populations are more complex and heterogenous than previously thought ^18^, and recent scRNA-seq analysis showed that a pericyte subpopulation promoted colorectal cancer metastasis ^24^.

Interactions between pericytes and other TME cells are still unknown, largely due to the limited availability of gene signatures that identify pericytes compared to other TME cell types in patient-derived cancer samples. We aimed to study pericytes profile across cancers and their communications with immune cells, other stromal cells, and tumor cells. Therefore, we adopted the implementation of a deeper analysis method of transcriptomic data sets from different tumor samples. The emergence of single-cell profiling technologies allows the measurement of the gene expression profiles of individual tumor, immune, and stromal cells in a high throughput fashion to determine their role in cancers ^25^. Single-cell technologies have improved our understanding of cancer biology, providing insights into tumor heterogeneity, TME cell interactions, and treatment resistance mechanisms ^26–29^. We performed a pan-cancer data analysis on: colon, skin, breast, bladder, pancreatic, ovarian, prostate, head and neck, bile duct, and cervical cancers, neurofibroma (benign tumor), lymphomas and lung as well as PTC and ATC, to characterize pericyte enrichment and expression profiles in the TME. For pericytes characterization, we have used research approach that analyzes and combines multiple scRNA-seq datasets from different studies to draw broader biological conclusions. This technique enhances statistical power, improves reproducibility, and can uncover patterns that might not be evident in individual studies. Our characterization identified a distinct pericyte gene signature as compared to fibroblasts in different cancers, including thyroid cancers (PTC and ATC). Furthermore, the cell-to-cell communication analysis revealed that pericytes are the most active cells (highest ligand secretion) in the TME and interact with tumor cells, e.g. with immune T cells in thyroid cancer. Pericytes interaction with T cells may cause T-cell exhaustion, thus effectively impairing their ability to eliminate tumor cells. Moreover, the single-cell data analysis across other cancers enabled the identification of pan-cancer pericytes signature.

We have also investigated the prognostic relevance of pericyte signatures from thyroid cancer and other life-threatening cancers providing new biological and translational insights through survival analysis. The associations of pericytes levels in TME and its impact on patient survival was assessed in different cancer data sets from The Cancer Genome Atlas Program (TCGA) ^30–39^. We hypothesize that pericytes may cause an immunosuppression in the TME, leading to tumor progression and poor over-all survival.

In summary, our study, for the first time, provides a pan-cancer analysis that identifies a universal pericyte signature associated with poor clinical outcomes. This provides a potential novel robust multiplex biomarker for the early identification of cancer. Ultimately, pericytes specific genes identified in this study could be targeted, providing innovative therapeutic strategies tailored to the TME.

## MATERIALS AND METHODS

### Datasets analysis

To perform scRNA-seq analysis in thyroid cancer, we used publicly available data from the Gene Expression Omnibus (GEO) database, specifically dataset GSE193581 ^27^. This dataset comprises raw single-cell data from patients with PTC (n=7), ATC (n=10), and normal thyroid (NT) (n=6) samples. The authors of the original manuscript focused their analyses on thyroid tumor cells and immune cells, while largely analyzing the fibroblast stromal cell populations ^27^. Notably, our prior work demonstrated that pericytes contribute to drug resistance in thyroid cancer models ^13–15^, which further emphasized the importance of characterizing pericytes at single-cell level in patient samples to decipher their role in thyroid cancer progression ^13,14^. This study was performed using publicly available human samples datasets **(Supp. Table 3)** without any patient healthcare information, so it does not require Institutional Review Board Committee (IRB) approval.

### Single-cell transcriptomics data analysis

The raw gene-cell count matrices from each sample were analyzed using Python 3.11, leveraging the Scanpy library and other supportive Python packages ^40^. Low-quality cells were filtered using Scanpy 1.10, retaining only cells with more than 100 unique genes and less than 30% mitochondrial Unique Molecular Identifiers (UMI’s). Genes expressed in fewer than three cells were excluded from the analysis to ensure robust downstream processing. The count matrices were normalized using Scanpy’s default method, which scales each cell’s total counts across all genes to a uniform value, ensuring comparable cell-to-cell data ^40^. This normalized data was subsequently log-transformed to mitigate random variance and highlight meaningful biological differences. The processed data was then used to identify 2,000 highly variable genes using scanpy.pp.highly_variable_genes function ^41^, which were subjected to principal component analysis (PCA). PCA reduced the dimensionality of the dataset by identifying principal components that captured the most variance across samples. To cluster cells with similar profiles, Leiden clustering was performed on the top principal components using Scanpy ^41^. This analysis was visualized using Uniform Manifold Approximation and Projection (UMAP), providing a two-dimensional map that revealed relationships among cell populations.

### Cell types and pericytes annotation

The cell clusters were annotated to cell types using automatic and manual annotation approaches. Canonical cell markers from previous studies guided manual annotation ^27^, while automatic annotation was performed using Celltypist with the Immune Cell Database ^42^. Notably, neither the automatic nor the manual annotation successfully identified pericytes within the dataset. To overcome this limitation, we conducted a more refined annotation based on genes known to be highly expressed or minimally/non-expressed in pericytes, as reported in our previous study **(Table 1)** ^13^. These genes were identified through a comprehensive PubMed literature search and subsequently used to annotate pericytes. A pericyte gene set score was then computed for each cell using using *scanpy.tl.score_genes* function in the Scanpy tool ^40^, enabling quantitative assessment of pericyte-like transcriptional profiles.

**Table 1.**
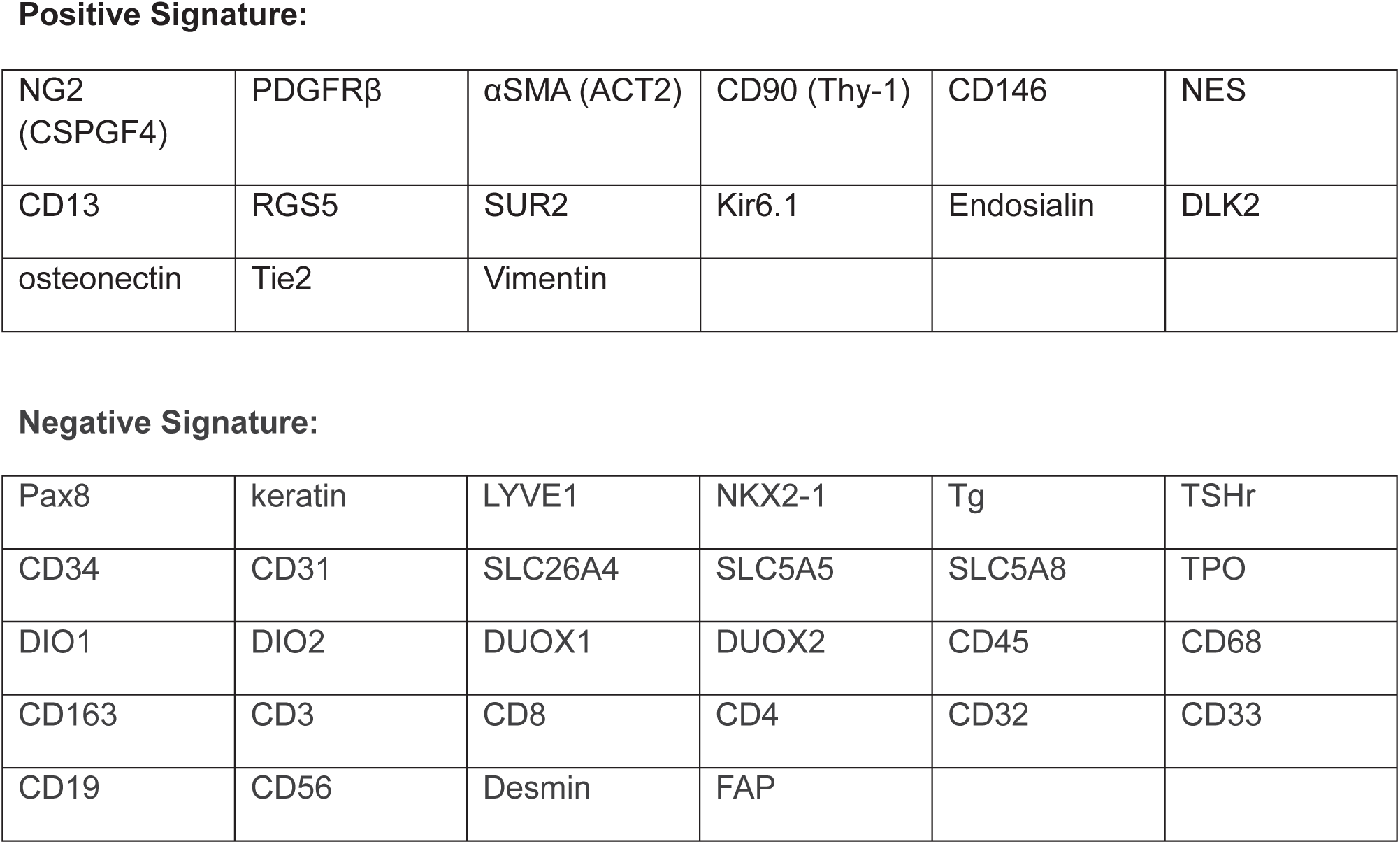
Pericyte gene signature by Iesato A. et al [PMID: 34302727] used for annotation of the pericyte cell type in single-cell data.

**Positive Signature:**
|  |  |  |  |  |  |
| --- | --- | --- | --- | --- | --- |
| NG2<br>(CSPGF4) | PDGFRβ | αSMA (ACT2) | CD90 (Thy-1) | CD146 | NES |
| CD13 | RGS5 | SUR2 | Kir6.1 | Endosialin | DLK2 |
| osteonectin | Tie2 | Vimentin |  |  |  |
| **Negative Signature:** |  |  |  |  |  |
| Pax8 | keratin | LYVE1 | NKX2-1 | Tg | TSHr |
| CD34 | CD31 | SLC26A4 | SLC5A5 | SLC5A8 | TPO |
| DIO1 | DIO2 | DUOX1 | DUOX2 | CD45 | CD68 |
| CD163 | CD3 | CD8 | CD4 | CD32 | CD33 |
| CD19 | CD56 | Desmin | FAP |  |  |

### Identification of cell type signatures

After cell type annotation, we identified genes that were significantly over-expressed in the pericytes as compared to fibroblasts in the dataset. Fibroblasts were annotated based on an automatic tool CellTypist gene signature **(Table 2)** ^42^. The significantly over-expressed genes were identified by implementing the Wilcoxon Rank Sum test statistical test and fold change. The genes that achieved adjusted p<0.05 and average log2 fold change (FC)> 0.25 were considered significantly over-expressed in pericytes.

**Table 2.** Gene signature used for fibroblast annotation in single-cell data.

|  |  |  |  |  |  |
| --- | --- | --- | --- | --- | --- |
| COL1A1 | COL1A2 | DCN | SMOC2 | SFRP1 | NTRK2 |
| PRRX1 | EBF2 | OLFML1 | MXRA5 | F10 | ANGPTL1 |
| PDGFRA |  |  |  |  |  |

### Cellular communication analysis

To investigate cell-cell communication, particularly involving pericytes, we utilized the CellChat platform ^43^. Pericytes, fibroblasts, tumor and immune cells were subsetted from along with ATC tumor cells from GSE193581 ^27^, and ligand-receptor (L-R) analysis was performed using the standard CellChat pipeline ^43^. Differentially expressed signaling genes were identified through the Wilcoxon rank-sum test (p<0.05), followed by the calculation of communication probability and interaction strength between cell types. To compare the overall signaling architecture between cell types, we analyzed interaction weights, which represent the aggregate information flow of all L-R interactions between specific cell types, such as pericytes and T-cell lineages. The interaction weights quantify the likelihood of cell-cell communication occurring between cell types through specific signaling pathways. For example, cells with high expression of a ligand are more likely to exhibit high interaction scores with cells expressing the corresponding receptor.

### Pericytes enrichment-based patient survival estimation

To explore association of top 50 thyroid cancer over-expressed genes from pericytes signature **(Suppl. Table 2, Fig.2G)** with overall survival, we performed survival association analysis. The survival analysis for pericytes was conducted using the PTC dataset from TCGA, comprising nearly 500 PTC samples ^35^. The pericytes score within the TCGA samples was calculated based on expression of top 50 pericytes genes **(Suppl. Table 2, Fig.2G)** using the Gene Set Variation Analysis (GSVA) approach ^44^. These scores were stratified into high and low groups by determining the optimal cut point via the Cutpoint Algorithm ^45^. Survival analysis was subsequently conducted using the Survival Genie platform ^46^, which allows the assessment of pericyte scores with favorable or poor survival in patients with cancer.

### Pericytes characterization in other cancers and pericytes signatures generation

To further characterize pericytes in other cancers, we obtained and analyzed the dataset from Gene Expression Omnibus (GEO) or Tumour Immune Single-cell Hub ^47^ databases. The raw data for each cancer ^48–68^ were processed to remove low quality cells, normalized, and clustered using the Seurat package V5 ^69^. The transcriptional clusters were annotated based on immune cell markers as well as calculating pericyte score using our previously published pericyte gene signature ^13^ **(Table 1).** If we detected a cluster/s in the dataset with high module score for pericyte genes, we annotated it as pericytes. We performd this single-cell analysis on all datasets (n=22) from 21 studies ^48–68^ corresponding to cancers and an additional dataset from a benign tumor (neurofibroma) ^48^ **(Suppl. Table 3)**. After final annotation of each dataset, we generated transcriptional signatures of pericytes from each dataset using differential gene expression analysis (DEGs). We used the Receiver Operating Characteristic (ROC)-based DEGs analysis in Seurat’s FindMarkers ^69^ to identify genes highly expressed in pericytes as compared to fibroblasts. Genes with an AUC >0.6 were selected as potential pericyte over-expressed markers based on their ability to discriminate pericytes from fibroblasts.

### Universal pericytes signature

To develop a universal pericyte signature, we integrated gene expression data across multiple individual datasets analyzed in the previous section ^27,48–68^. Genes that were consistently overexpressed in pericytes in more than 60% of the datasets were selected. These genes were ranked based on both the frequency of over-expression and their average fold change across datasets. To ensure specificity and exclude genes expressed in normal immune cells, we analyzed the 10X Genomics legacy peripheral blood mononuclear cells (PBMC) 10K dataset ^69,70^. This step ensures that genes are expressed by pericytes but not by normal immune cells reducing the toxic effects if targeted in future therapy development. Genes with an absolute RNA expression value greater than 0.1 in any immune cell type were filtered out. The PBMC dataset was processed through normalization, clustering, and cell-type annotation before mapping the expression of candidate pericyte genes using Seurat workflow ^69^.

### Cancer clinical outcome associated with pericyte gene signature

To assess the cancer prognostic relevance of the universal pericyte gene signature, we performed association with overall survival using the Survival Genie platform across the following TCGA datasets: bladder, breast, ovarian, pancreatic, prostate, colorectal, cholangio, cervical, head and neck, and thyroid (including PTC) carcinomas, as well as melanoma, lung adenocarcinoma and lung squamous cell carcinoma ^30–37,39,71–73^. These cancer types were chosen as we have analyzed their single-cell datasets to generate pericyte signatures. Survival analysis was conducted using the Cox proportional hazards model, stratifying gene expression into high and low groups based on an optimal cut-point algorithm ^46^. To identify a subset of the universal pericyte signature associated with poor clinical outcomes, we selected genes that demonstrated a statistically significant association with poor survival in at least 30% of the cancer datasets. These outcome-associated genes were then ranked based on both the magnitude of overexpression and the strength of their association with survival outcomes (HR values) to generate a refined pan-cancer pericyte signature.

### Pathways and interactive network analysis to achieve mechanistic insights

To gain mechanistic insights, we performed pathway enrichment analysis using the Enrichr platform ^74^ on genes associated with poor survival across different cancer types. This platform integrates multiple pathway databases to assess the over-representation of input gene list. In this study, enrichment was specifically carried out using the Reactome and Cancer Hallmark pathway databases^75,76^. Pathways with a Benjamini–Hochberg corrected p-value <0.05 were considered significantly affected. We also performed interactive network analysis based on co-expression, physical interactions, text mining, and shared biological processes using the GeneMANIA plugin in Cytoscape platform ^77,78^. Networks were visualized using Cytoscape platform and analyzed using network analyzer package. Master regulators within the network were identified based on their degree of interaction within the network.

### Correlation of cancer-associated pericyte signature and T cell exhaustion genes

To investigate the correlation between pericyte-associated genes and T cell exhaustion markers, we analyzed pan-cancer normalized bulk RNA-sequencing data downloaded from the cBioPortal platform ^79^. The dataset included expression profiles from 10,967 samples across 32 cancer studies **(Suppl. Table 7)**, all part of the TCGAs pan-cancer collection ^38^. We specifically assessed the correlation between pan-cancer pericyte genes signature and established exhaustion markers such as HAVCR2 (also known as TIM-3). Spearman rank correlation analysis was performed in R, and statistical significance was determined using the cor.test function ^80^. Genes with a Spearman R²>0.10 and a p-value <0.001 were considered statistically significant associated with immune T cell exhaustion.

### Scripts and workflow

Well-developed and published programs such as Scanpy v1.11.4 ^40^, CellTypist v1.7.1 ^42^, Seurat v5.3 ^69^, CellChat v2.x ^43^, GSVA ^44^, Survival Genie 2.0 ^46^, Enrichr ^74^, and Cytoscape v3.10.3 ^78^ with the GeneMANIA ^77^ and GEO datasets ^48–68^ have been used for the analyses. All analyses were conducted using Python/R workflows and scripts, developed with standardized tools for single-cell analysis.

## Results

Single-cell profiling enables high-resolution dissection of tumor heterogeneity, allowing us to define the specific roles of tumor, stromal, and immune cell populations in cancer initiation and progression. Among the stromal cells, pericytes, which wrap around blood vessels, emerged as a key cell type of interest due to their suggested role in regulating angiogenesis and tumor progression^18^. While we have previously characterized pericytes in bulk tissue samples from human thyroid cancer and mouse models ^13,14^, their precise molecular identity and functional roles at single-cell resolution remain poorly defined.

To address this gap, we designed a comprehensive study aimed at characterizing the single-cell profile of pericytes across different cancers. An overview of our approach is shown in **Fig.1**. We began by analyzing pericytes from single-cell RNA-sequencing dataset of NT, PTC, and ATC ^27^, involving rigorous data preprocessing, accurate pericyte annotation, and identification of their gene expression signatures. We then conducted cell-cell communication analyses to understand their interactions within the TME **(Fig.1A)**. Following the thyroid cancer analysis, we expanded our study to additional cancer types using 22 publicly available single-cell datasets from 21 studies^48–68^ **(Suppl. Table 3**, **Fig.1B)**. This allowed us to generate a universal pericyte gene signature composed of genes consistently over-expressed by pericytes across different cancers and benign tumors (i.e. neurofibroma) **(Fig.1B)**.

**Figure 1.**
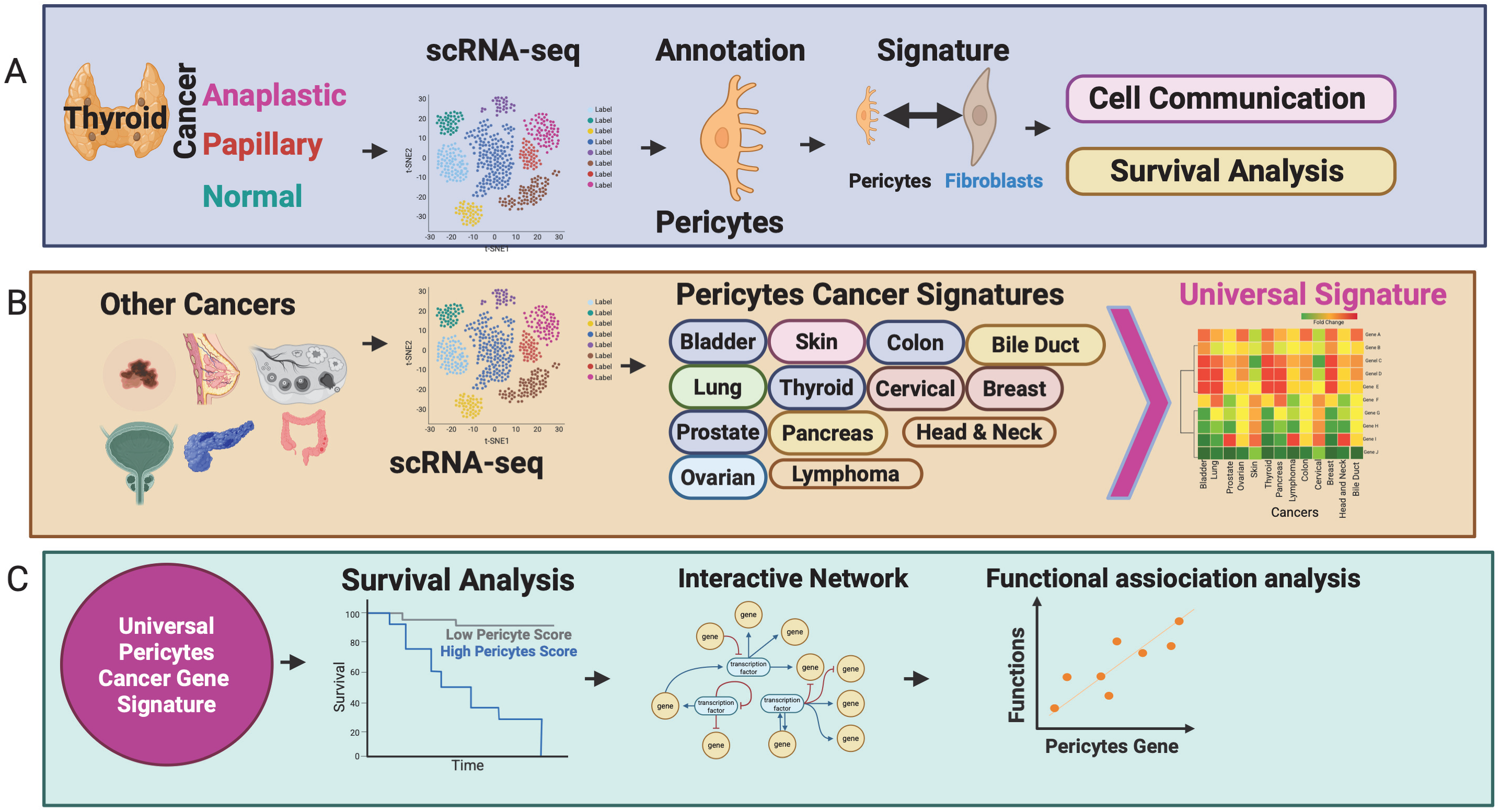
An overview of a study investigating the association between pericytes and cancer outcomes using a comprehensive analytical framework. **A.** schematic overview of the single-cell analysis pipeline used to annotate, characterize, and determine the clinical significance of pericytes in thyroid cancer. **B.** Extension of this analysis to other cancer types, including breast, ovarian, bladder, skin, colorectal cancers, etc. to evaluate the broader biological relevance of pericytes. Comparative analysis across cancer types led to the identification of a universal pericytes signature. **C.** The biological significance and prognostic impact of this signature were further explored through interactive network modelling, survival association analysis, and correlation with immune phenotypes.

We further refined this signature by integrating survival association data to identify genes correlated with poor clinical outcomes that generated the first pericytes cancer-associated gene signature. To gain mechanistic insights, this signature was evaluated by pathways and interactive network analysis **(Fig.1C)**. This study presents the first universal pericyte gene signature linked to cancer prognosis, offering novel insights into pericyte biology and providing a foundation for developing pericyte-targeted therapeutic strategies in cancer.

### Characterization of pericytes gene signature in thyroid cancer

The analysis of thyroid cancer single-cell genomics data ^27^ after normalization and filtering out low-quality cells identified 28,890 cells for downstream analysis. Using PCA based on 2,000 highly variable genes and clustering with the Leiden algorithm, we identified 40 distinct cell clusters **(Fig.2A)**. Some clusters were composed exclusively of cells from ATC or PTC samples, without overlap with NT cells profiles, indicating cancer-specific clusters (**Suppl. Table 4**). In contrast, clusters representing immune or stromal cell types contained cells from NT, ATC, and PTC samples, suggesting shared transcriptional profiles **(Suppl.Fig.1A, Suppl. Table 4)**. To identify the cell types, we performed both manual **(Fig.2B)** and automatic (using Cell Typist) **(Fig.2C)** annotation, leveraging cell-specific markers from an original study ^27^. This analysis successfully identified immune and stromal cell populations, including T cells, B cells, fibroblasts, but not pericytes **(Fig.2B-C, Suppl.Fig.1B**, **Table 2)**. Manual annotation for single-cell analysis is often more accurate than automatic annotation because it leverages expert knowledge to interpret complex or ambiguous gene expression patterns that algorithms might misclassify. It also allows for greater flexibility and oversight, especially when dealing with novel, rare cell types, or transitional cell states that automated tools may not be trained to recognize ^81^. Automatic annotation in single-cell analysis can make errors because it relies heavily on reference datasets and predefined marker genes, which may not fully capture the diversity or context-specific expression patterns of cell types in new datasets ^81^. If the reference is incomplete, biased, or not representative of the sample being analyzed, the algorithm may misclassify cells. Therefore, we performed an additional analysis using our previously defined pericyte associated genes ^13^ **(Table 1)** by calculating the average expression and subtracting the expression of negatively associated genes **(Table 1)**. This highlighted cluster 18 **(Fig.2D-E)** due to is high expression of pericyte genes **(Table 1)**. All these processing steps culminated in the final cell-type labeling, including pericyte cluster **(Fig.2F)** through manual annotation based on the markers shown in **Fig.2B**, along with **Fig.2D-E** analysis.

**Figure 2.**
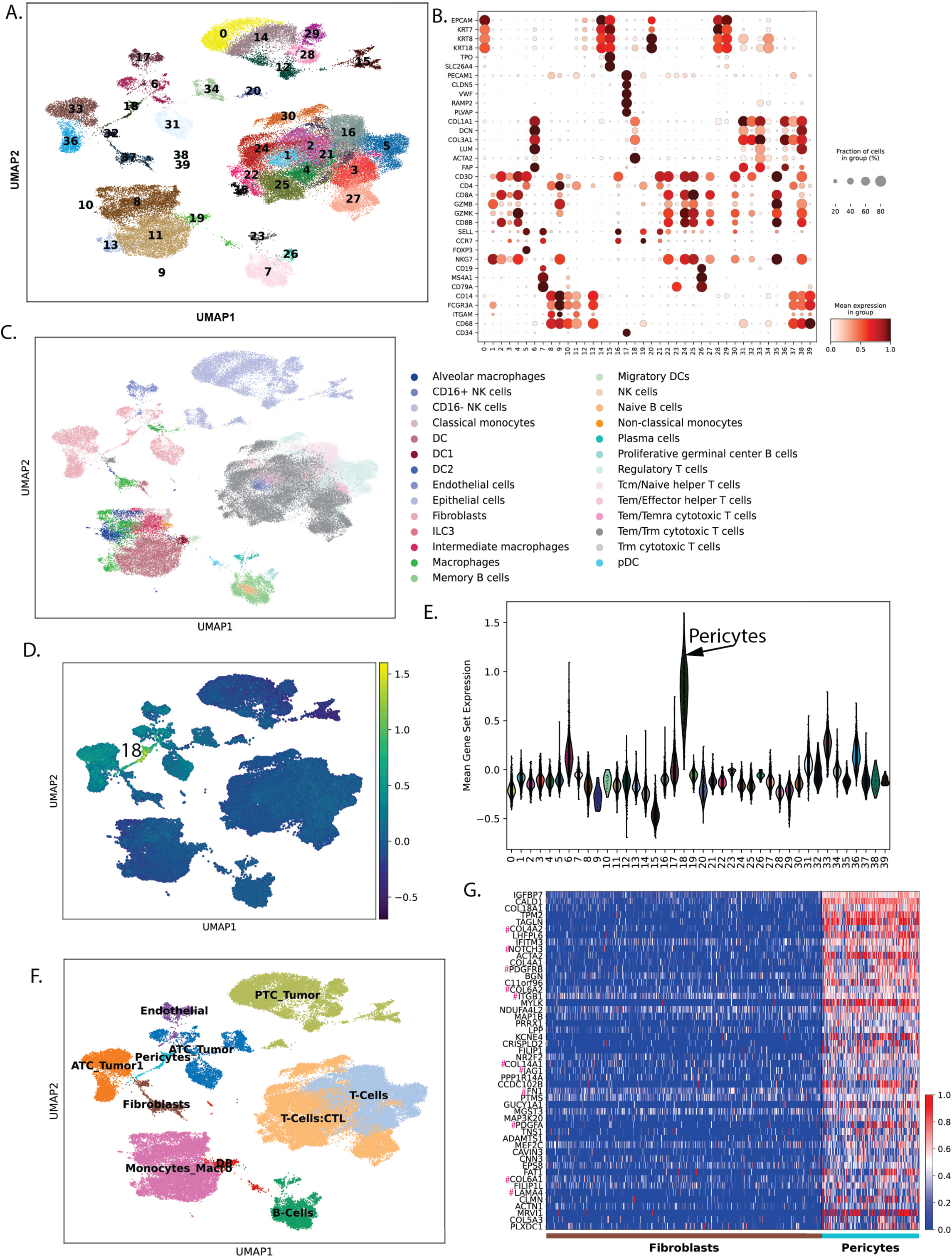
Comprehensive characterization of pericyte populations in thyroid cancer. **A.** UMAP plot showing the distribution of 40 transcriptionally distinct clusters derived from single-cell RNA sequencing data of thyroid cancer and adjacent normal tissues analysis. These clusters were identified through unsupervised clustering and represent diverse cellular populations present within the tumor microenvironment and normal thyroid tissue **B.** Dot plot representing the expression patterns of well-established canonical marker genes across the 40 clusters, enabling preliminary identification of major cell types including epithelial cells, immune cells, stromal cells, and vascular-associated populations. **C.** Automated annotation of cell identities using the CellTypist tool, which classified the clusters into granular cell categories. **D.** UMAP feature plot visualizing the distribution of a pericyte score computed for each cell based on the expression of a curated pericyte-specific gene signature (Table 1). This highlights pericyte-enriched clusters within the dataset. **E.** Violin plot displaying the distribution of pericyte scores across all 40 clusters, revealing cluster 18 with elevated scores suggestive of pericyte existence in the dataset (0 = PTC_Tumor, 1 = T-Cells:CTL, 2 = T-Cells, 3 = T-Cells, 4 = T-Cells:CTL, 5 = T-Cells, 6 = ATC_Tumor, 7 = B-Cells, 8 = Monocytes_Macro, 9 = Monocytes_Macro, 10 = Monocytes_Macro, 11 = Monocytes_Macro, 12 = PTC_Tumor, 13 = Monocytes_Macro, 14 = PTC_Tumor, 15 = PTC_Tumor, 16 = T-Cells, 17 = Endothelial, 18 = Pericytes, 19 = DB, 20 = PTC_Tumor, 21 = T-Cells, 22 = T-Cells:CTL, 23 = B-Cells, 24 = T-Cells:CTL, 25 = T-Cells:CTL, 26 = B-Cells, 27 = T-Cells:CTL, 28 = PTC_Tumor, 29 = PTC_Tumor, 30 = T-Cells:CTL, 31 = ATC_Tumor, 32 = Fibroblasts, 33 = ATC_Tumor1, 34 = ATC_Tumor, 35 = T-Cells:CTL, 36 = ATC_Tumor1, 37 = Fibroblasts, 38 = DB, 39 = Monocytes_Macro). **F.** UMAP plot showing the final classification of major cell types obtained by integrating automated annotation with manual expert curation. **G.** Heatmap showing the top genes significantly overexpressed in pericytes compared to fibroblasts (based on relative expression) within the thyroid cancer dataset, highlighting molecular features that distinguish these two closely related stromal populations. Note: # indicates those genes with functions of ligand or receptor within the pericyte signature.

After annotation, a differential gene expression signature for pericytes was generated by comparing their transcriptional profile with that of fibroblasts **(Fig.2G, Suppl. Table 1)**. The pericytes over-expressed 374 genes **(Suppl. Table 1)** were identified from the profile based on fold change and BH-adjusted p-value (p<0.05) using the Wilcoxon Rank sum test. The heatmap of the top 50 over-expressed genes **(Fig.2G, Suppl. Table 2)** in pericytes based on Scanpy ^40^ generated score from scanpy.tl.score_genes function depicts a clear segregation of pericytes compared to fibroblasts in thyroid cancer (GSE193581 dataset). Based on our knowledge, this is the first time anyone has characterized thyroid cancer-enriched single-cell pericytes in clinical samples. The pericytes formed a single cluster (i.e., cluster ID=18), independent of their origin from ATC or PTC, suggesting a shared molecular profile.

### Pericytes strongly interact with tumor cells and T-cells, and are associated with poor survival

Following annotation of pericytes and the generation of gene signatures, we conducted an innovative analysis to uncover how these cells communicate with tumor cells and other components of the TME. Using the CellChat platform^43^, we analyzed cell-cell communications, assessing ligand-receptor interactions and their correlation across different cell types. This approach revealed that pericytes engage in extensive crosstalk with tumor cells and immune cells **(Fig.3A)**. The pericytes show highest interactions with T-cells and tumor cells **(Fig.3A)**. The pericytes are the top signal senders after tumor cells, indicating their involvement in shaping the TME **(Fig.3B)**. On the other hand, T-cells are the top signal receiver along with endothelial and fibroblasts **(Fig.3B).** Further examination of the pathways that are secreted by pericytes includes Collagen, MHC-1, Fibronectin 1, and Angiopoietin, etc. COL4A1 on pericytes interacts with CD44 on T cells, potentially hindering T cell migration, activation, and adhesion within the extracellular matrix (ECM) to eradicate tumors **(Fig.3 C-D)**. Additionally, pericytes Nectin-2 binds to TIGIT on a T cell **(Fig.3C-D)**, preventing the T cell from effectively attacking tumor cells ^82^. Collectively, these novel insights highlight a possible mechanism by which pericytes contribute to immune escape.

**Figure 3.**
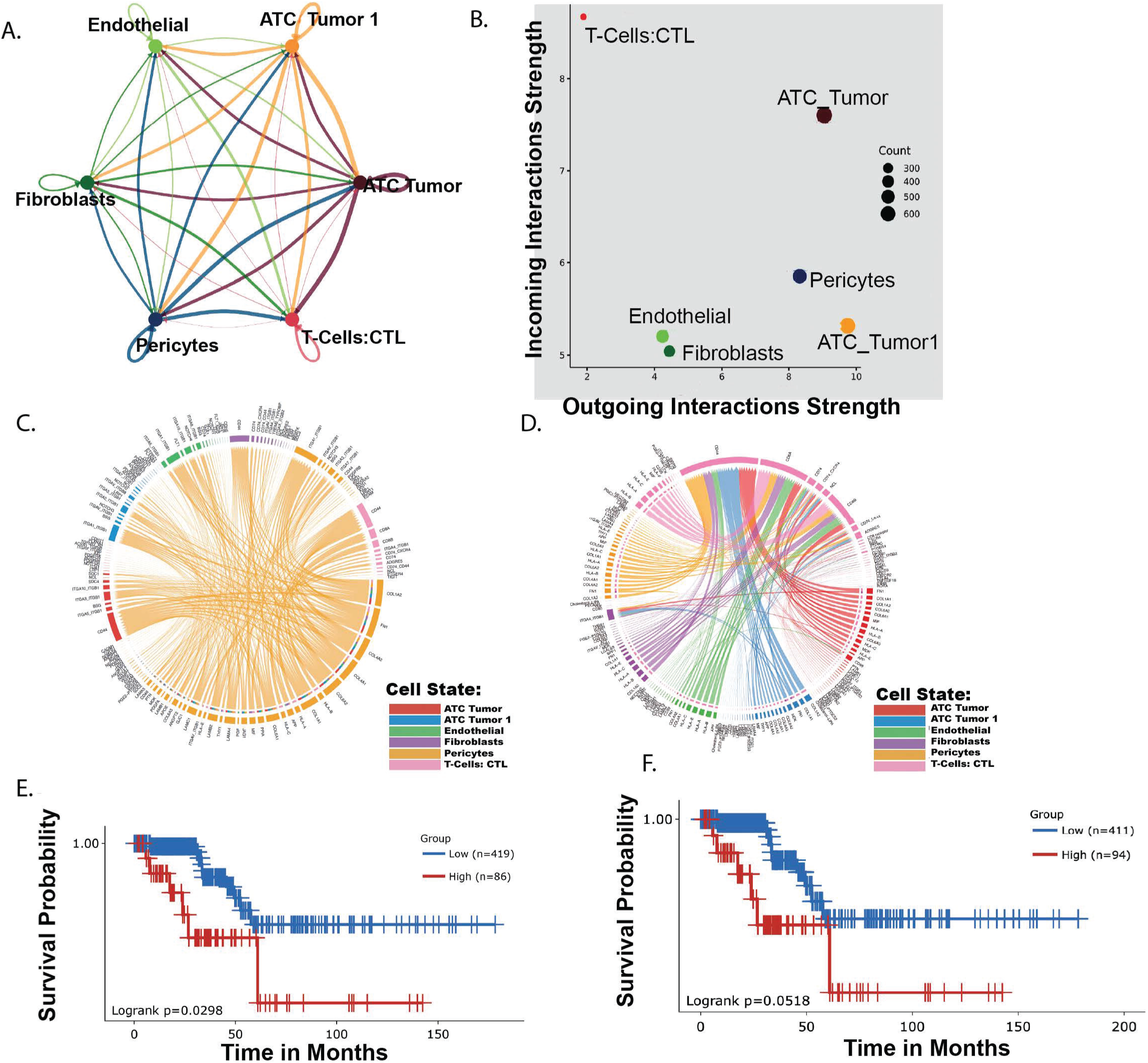
Cellular communication and survival associations of pericytes. **A.** Cellular communication network depicting the signal strength of interactions among tumor cells, pericytes, fibroblasts, and immune cells. Edges represent communication interactions, with different colors indicating the cell type of origin and edge width reflecting interaction strength. **B.** Scatter plot showing sum of incoming and outgoing interaction numbers across cell types. The data reveal that pericytes exhibit the highest outgoing interactions, comparable to tumor cells. **C.** Circos plot showing the communication landscape of pericytes with other cell types: e.g. pericytes interact with CD8+ T cells through MHC-I:CD8 and collagen:CD44/integrin-mediated signalling axes. **D.** Circos plot summarizing communication from all cell types, with edge colours representing the origin (source) cell type. **E.** Kaplan–Meier survival analysis based on the top 50 genes over-expressed in pericytes compared to fibroblasts, using TCGA papillary thyroid cancer (PTC) datasets. Patients were stratified into high and low expression groups based on the enrichment of pericyte-associated gene signatures using optimal cut-point analysis. Elevated expression of pericyte-enriched genes was associated with significantly worse survival outcomes (shown with red line). **F.** Similar survival analysis as in panel E but restricted to ligand–receptor gene pairs expressed in the pericyte signature (marked by # in Fig.2G). The analysis was conducted using the Survival Genie platform.

To further support the role of pericytes in thyroid cancer prognosis, we conducted a survival analysis using RNA-seq data of PTC samples (n=505) profiled in the TCGA initiative ^35^. In this dataset, the overall survival analysis was performed using the top 50 pericyte-expressed genes shown in **Fig.2G**. The analysis depicted samples with higher expression of pericyte genes have a poorer outcomes (HR=2.9, p-value=0.02), suggesting that the pericyte gene signature is associated with overall poor survival in PTC patients **(Fig.3E)**. Given our hypothesis that communication between pericytes and immune cells is critical for supporting thyroid cancer progression, we specifically focused on the overall survival analysis of ligands and receptors identified within the pericyte signature **(Fig.2G**, marked by #). Our analysis revealed that the enrichment of pericyte-derived ligands and receptors also correlates with poor survival outcome in PTC patients (HR=2.6, p-value=0.05, **Fig.3F)**. In contrast, patients with lower expression levels of these pericyte-associated signaling molecules demonstrated better survival rates **(Fig.3F)**.

These findings provide evidence that pericytes play a pivotal role in promoting poor outcomes in thyroid cancer by facilitating immune escape and supporting tumor growth.

### Pan-cancer pericytes characterization from single cell data

To further extend our study of pericytes and characterize their presence across diverse cancer types, we conducted an analysis of single-cell RNA-sequencing from 23 datasets from 21 studies corresponding to 15 cancers and 1 benign tumor (neurofibroma): breast, pancreatic, bladder, ovarian, non-small cell lung cancer, bladder, cervical, prostate, basal cell, colon, head and necks cancers, melanoma, lymphomas, as well as thyroid cancers (PTC and ATC) ^27,48–68^ **(Suppl. Table 3)**. Each dataset underwent quality control, normalization, clustering, and cell-type annotation using canonical lineage markers ^69^. During annotation, we applied the expression of a curated pericyte gene signature **(Table 1)** to annotate clusters that depict high expression of pericytes gene set. For example, in colon cancer, we identified 29 clusters **(Fig.4A)** that were categorized into different cell types **(Fig.4D)** based on marker expression **(Fig.4B)** and pericyte signature enrichment **(Fig.4C)**. Similar analysis was done on datasets from other tumors including: melanoma **(Fig.4E, Suppl.Fig.2-3)**, breast **(Fig.4F, Suppl.Fig.4)**, bladder **(Fig.4G, Suppl.Fig.5)**, pancreatic **(Fig.4H, Suppl.Fig.6-7)**, ovarian **(Fig.4I, Suppl.Fig.8-11)**, prostate **(Fig.4J, Suppl.Fig.12-14)**, head and neck **(Fig.4K, Suppl.Fig.15)**, cholangio-carcinoma **(Fig.4L, Suppl.Fig.16),** cervical **(Fig.4M, Suppl.Fig.17),** neurofibroma **(Fig.4N, Suppl.Fig.18)**, lymphoma **(Fig.4O, Suppl.Fig.19-20)**, lung **(Fig.4P, Suppl.Fig.21),** and thyroid **(Suppl.Fig.22)** tumor. Across the datasets analysed, we identified clusters with high score for pericytes signature genes **(Table 1)** as compared to fibroblasts annotated genes **(Table 2)**. After annotation in each dataset, we generated lists of genes significantly over-expressed (AUC ≥0.6 using *findmarkers* function in Seurat package ^69^) in pericytes compared to fibroblasts. This resulted in the generation of 22 distinct pericyte gene signatures spanning 15 cancer types and 1 benign tumor (neurofibroma). After including the pericyte signature from thyroid cancer dataset ^27^ **(Suppl. Table 2)** used for foundational analysis shown in **Fig.2**, we have in total 23 pericyte gene signatures.

**Figure 4.**
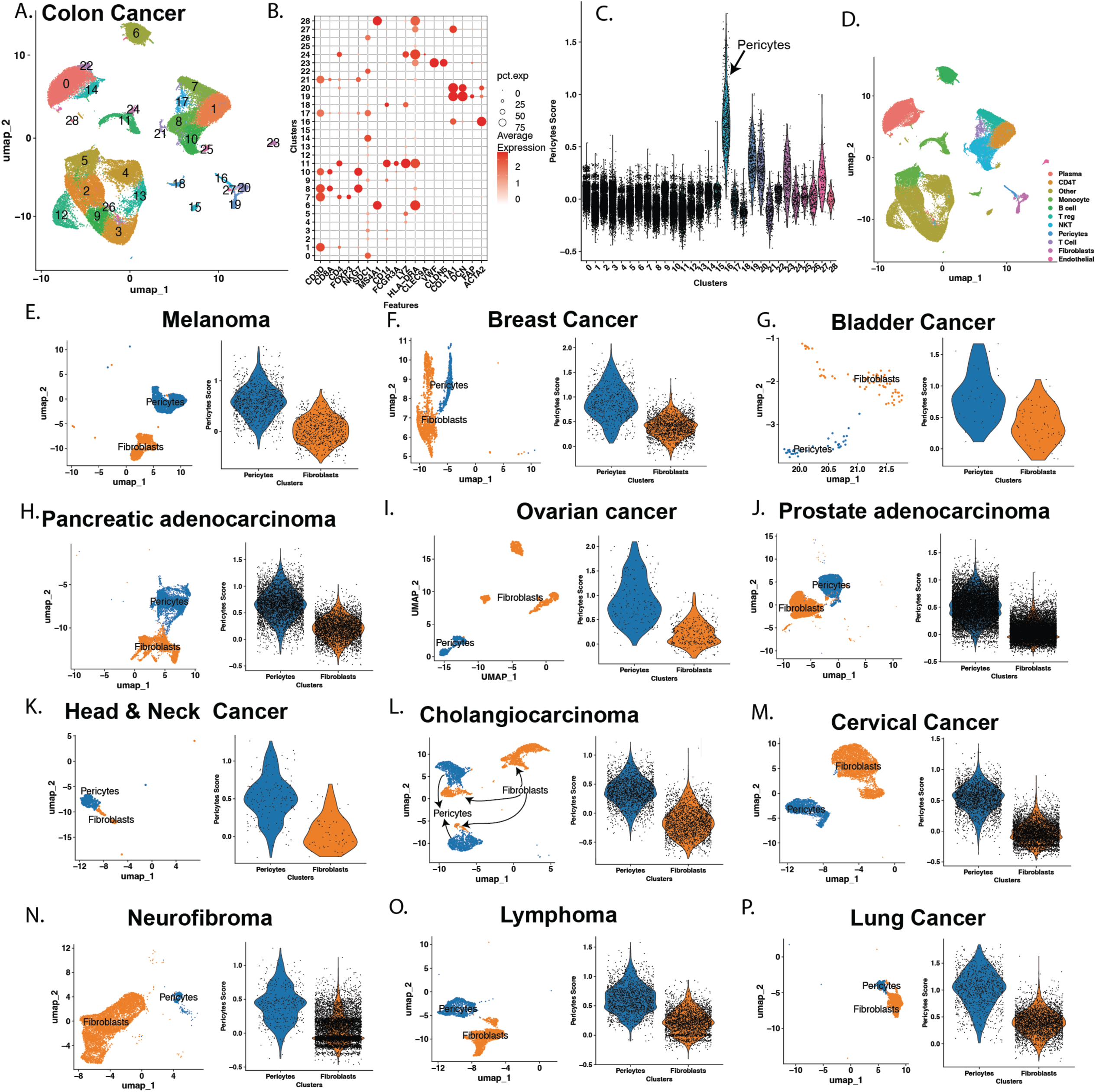
Pericytes enrichment analysis across different cancer types. **A.** UMAP visualization of pericyte profiling in colon cancer, using transcriptomic data obtained from the GEO or TISCH database. Clustering analysis identified multiple transcriptionally distinct clusters. 0=Plasma, 1=CD4T, 2=Other, 3=Other, 4=Other, 5=Monocyte, 6=B cell, 7=T reg, 8=NKT, 9=Other, 10=NKT, 11=Monocyte, 12=Other, 13=Other, 14=Plasma, 15=Other, 16=Pericytes, 17=T Cell, 18=Monocyte, 19=Fibroblasts, 20=Fibroblasts, 21=NKT, 22=Plasma, 23=Endothelial, 24=Monocyte, 25=Other, 26=Plasma, 27=Fibroblasts, and 28=Other. **B.** Dot plot showing the expression of marker genes used to annotate major cell types, including pericytes and fibroblasts, across the identified clusters in colon cancer. **C.** Violin plot depicting the distribution of pericyte scores across cell populations in colon cancer. This analysis supports the annotation of pericyte-enriched clusters based on pericyte-specific gene signatures (0=Plasma, 1=CD4T, 2=Other, 3=Other, 4=Other, 5=Monocyte, 6=B cell, 7=T reg, 8=NKT, 9=Other, 10=NKT, 11=Monocyte, 12=Other, 13=Other, 14=Plasma, 15=Other, 16=Pericytes, 17=T Cell, 18=Monocyte, 19=Fibroblasts, 20=Fibroblasts, 21=NKT, 22=Plasma, 23=Endothelial, 24=Monocyte, 25=Other, 26=Plasma, 27=Fibroblasts, and 28=Other). **D.** Final annotation of cell types in the colon cancer dataset. Similar analyses were performed across multiple other cancer types, including melanoma **(E)**, breast cancer **(F)**, bladder cancer **(G)**, pancreatic cancer **(H)**, ovarian cancer **(I)**, prostate adenocarcinoma **(J)**, head and neck squamous cell carcinoma **(K)** cholangiocarcinoma **(L)** cervical cancer **(M)**, neurofibroma (benign tumor) **(N)**, lymphoma **(O)**, and lung cancer **(P)**. For each of these cancers: Left panel: UMAP plot displaying cell clusters annotated as pericytes and fibroblasts based on transcriptional signatures. Right panel: Violin plot comparing the pericyte score between pericyte and fibroblast populations, showing an elevation of the score in pericyte-annotated clusters. This pan-cancer analysis demonstrates the consistent identification of pericyte populations across diverse tumor microenvironments and their distinction from fibroblasts based on transcriptional features.

### A comparative analysis across multiple cancer types identified a universal pericyte-associated gene signature

By integrating pericyte gene signatures from 22 cancer datasets and 1 benign tumor dataset (i.e. neurofibroma) **(Suppl. Table 3, Fig.5A)**, we identified a total of 1,648 genes significantly associated with pericyte in at least one dataset **(Fig.5B).** To ensure consistency, we filtered for genes expressed in ≥60% of these datasets, yielding a refined list of 100 genes (**Fig.5C**). *NOTCH3* emerged as the only gene consistently overexpressed across all pericyte signatures, followed closely by *ACTA2*, which was up-regulated in over 95% of the datasets **(Fig.5C)**.

**Figure 5.**
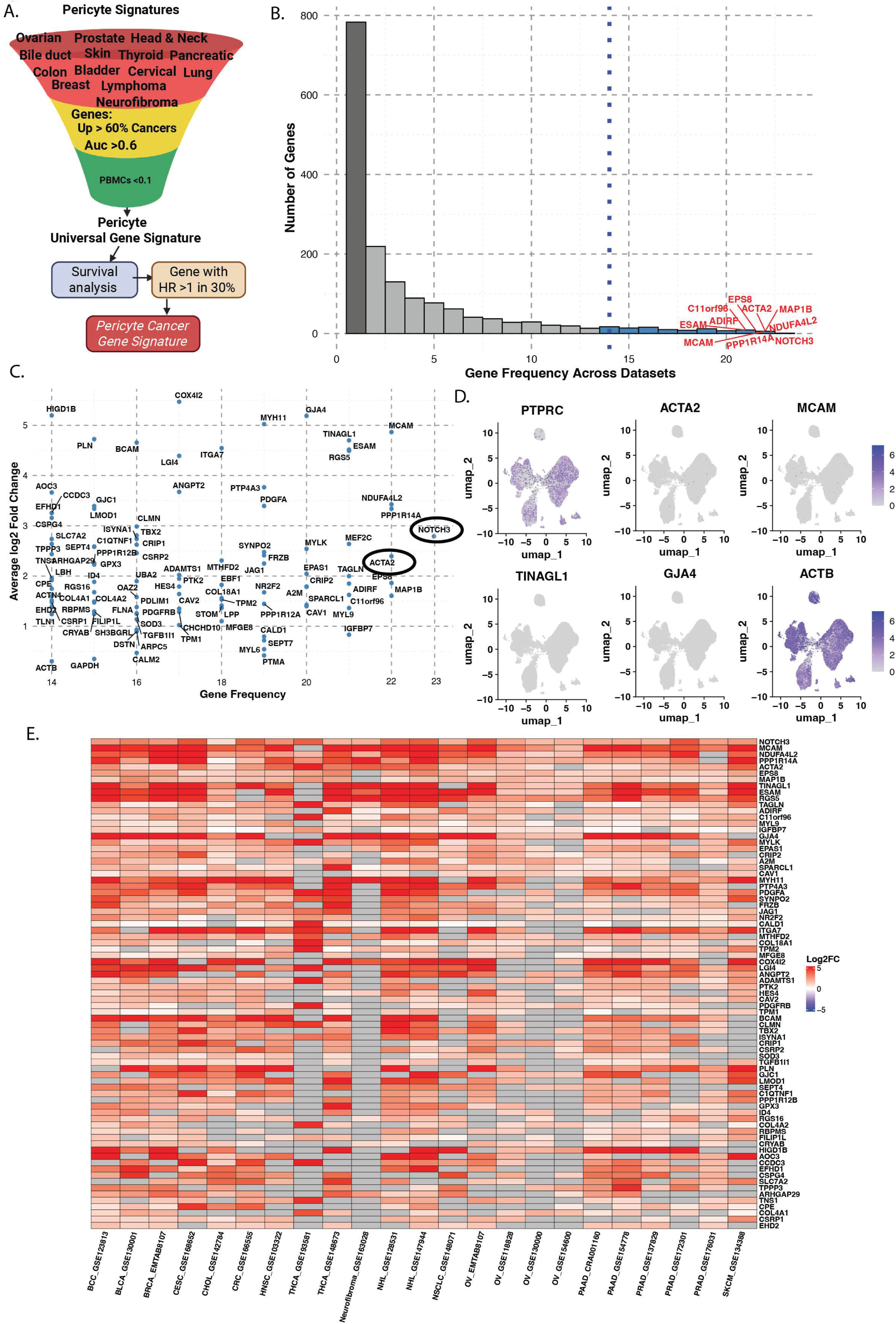
Universal pericyte gene signature from a pan-cancer analysis. **A.** Schematic overview of the steps to generate a pan-cancer–associated pericyte signature from single-cell datasets from different cancers. **B.** Binned histogram showing the frequency of genes over-expressed in pericytes across datasets. Genes over-expressed in more than 60% of datasets were considered candidate pericyte signature genes. The top 10 most frequent genes are highlighted in red. **C.** Scatter plot comparing the fold change and frequency of over-expression for pericyte-expressed genes that are over-expressed in pericytes from >60% datasets. **D.** Feature plots showing expression of selected pericyte signature genes in PBMC datasets from 10X Genomics, used as a control. PTPRC/CD45, a potent lymphocyte marker, is shown to be highly expressed in different Peripheral Blood Mononuclear Cells (PBMC) cell types (shown as positive control). In contrast, pericyte markers such as ACTA2 and MCAM show minimal expression in PBMCs, and housekeeping genes like ACTB are ubiquitously expressed and thus excluded from the final pericyte signature. **E.** Heatmap of the top 50 genes over-expressed in pericytes across more than 60% of datasets. Each row represents a gene, and each column represents a different single-cell dataset. The heatmap is pseudo-colored by log2 fold change, with red indicating high expression. Most of these genes are consistently over-expressed in pericytes across cancers, highlighting their universality.

To further enhance the specificity of this signature, we used 10x Genomics PBMCs datasets ^69,70^ to eliminate genes broadly expressed in immune populations. This filtering removed genes such as *MYL6* and *ACTB*, which showed high expression in T cells and monocytes **(Fig.5D)**. In datasets analyzed, pericytes have distinct cluster from cancer cells indicting pericytes distinct transcriptome profile and supporting association of these genes to pericytes. Through this systematic approach, we defined a core set of 78 genes **(Fig.5E**, **Suppl. Table 5)** that are robustly and selectively over-expressed in pericytes from all analyzed cancer dataset and minimally expressed in normal PBMCs cell types such as T-cells, myeloid cells and B cells. We refer to this refined set as the pericyte-universal signature, a putative pan-cancer pericyte-specific expression profile. The signature contains e.g. Melanoma Cell Adhesion Molecule (*MCAM*), which plays a role in pericytes and endothelial communication for regulating angiogenesis and blood barrier maintenance ^45,83^. The signature also includes Regulator of G-protein signaling 5 protein (*RGS5*), which is used for measuring pericyte coverage of vessels during development ^84^. *NOTCH3* is another pericyte-associated gene influencing pericyte differentiation and maturation ^85^. Additionally, signatures also have not-so-well-known pericytes-associated genes such as Caveolin-1 (*CAV1*), which is expressed in brain microvessels and might play a role in regulating permeability ^86^. Similarly, Protein Tyrosine Kinase 2 (*PTK2*) plays an important role in integrin and growth factor signaling that is essential for cell adhesion, migration, and mechano-transduction across many cell types ^24,87^.

### Pathways associations of the pericyte universal gene signature

Pathways linked to the over-expressed pericyte genes **(Fig.5E)** across different cancers play a pivotal role in tumor progression, immune modulation, and treatment resistance. Pathway enrichment analysis using the Reactome database revealed significant over-representation (BH-adjusted p<0.05) in signaling processes related to muscle contraction, RHO GTPase activity, Extracellular Matrix (ECM) organization, NOTCH3 activation, signaling by receptor tyrosine kinases, etc. **(Suppl. Table 6)**. Complementary analysis using the Cancer Hallmark database demonstrated significant enrichment (BH-adjusted p<0.05) in pathways associated with myogenesis, epithelial mesenchymal transition, angiogenesis, adipogenesis, NOTCH signaling, UV Response DN, and apoptosis. These pathways contribute to creating a tumor-supportive microenvironment by promoting cellular invasion, enhancing metastatic potential, tumor metabolism, and modulating immune cell infiltration. The conserved over-expression of these pericytes-related pathways across multiple cancer types highlights their potential as therapeutic targets and biomarkers for cancer prognosis and treatment stratification.

### Pan-cancer pericyte gene signature associated with poor outcomes

To investigate the clinical relevance of our previously identified universal pericyte gene set (n=78, **Suppl. Table 5**), we performed association analysis with overall survival using the Survival Genie platform across 13 cancer datasets from TCGA: TCGA-BLCA = bladder urothelial carcinoma ^71^; TCGA-BRCA = breast invasive carcinoma ^31^; TCGA-CESC = cervical squamous cell carcinoma and endocervical adenocarcinoma ^72^; TCGA-CHOL = cholangiocarcinoma ^88^; TCGA-COAD = colon adenocarcinoma ^30^; TCGA-HNSC = head and neck squamous cell carcinoma ^32^; TCGA-LUAD = lung adenocarcinoma ^36^; TCGA-LUSC = lung squamous cell carcinoma ^34^; TCGA-OV = ovarian serous cystadenocarcinoma ^33^; TCGA-PAAD = pancreatic adenocarcinoma ^39^; TCGA-PRAD = prostate adenocarcinoma ^37^; TCGA-SKCM = skin cutaneous melanoma ^73^; and TCGA-THCA = thyroid carcinoma ^35^. These datasets were selected based on the SC-RNA-sequencing studies that we analyzed to generate pericyte signatures **(Fig.4** and **Fig.2** for thyroid cancer). The analysis revealed that 19 genes were significantly associated with poor outcomes based on hazard ratio >1 and significant p-value (p<0.05) in at least 30% of the TCGA cancer data sets **(Fig.6A)**. Among them, TPM2 emerged as a top gene, linked to poor outcome in 9 out of 13 cancer types, followed by EHD2, which showed association in 7 datasets **(Fig.6B)**. These findings suggest that enrichment of pericyte-like gene expression **(Fig.6C)** may contribute to tumor resistance and poor prognosis. Network analysis of the 19 genes revealed a tightly connected interactome, indicating that these genes may act in concert to influence tumor progression **(Fig.6D)**.

**Figure 6.**
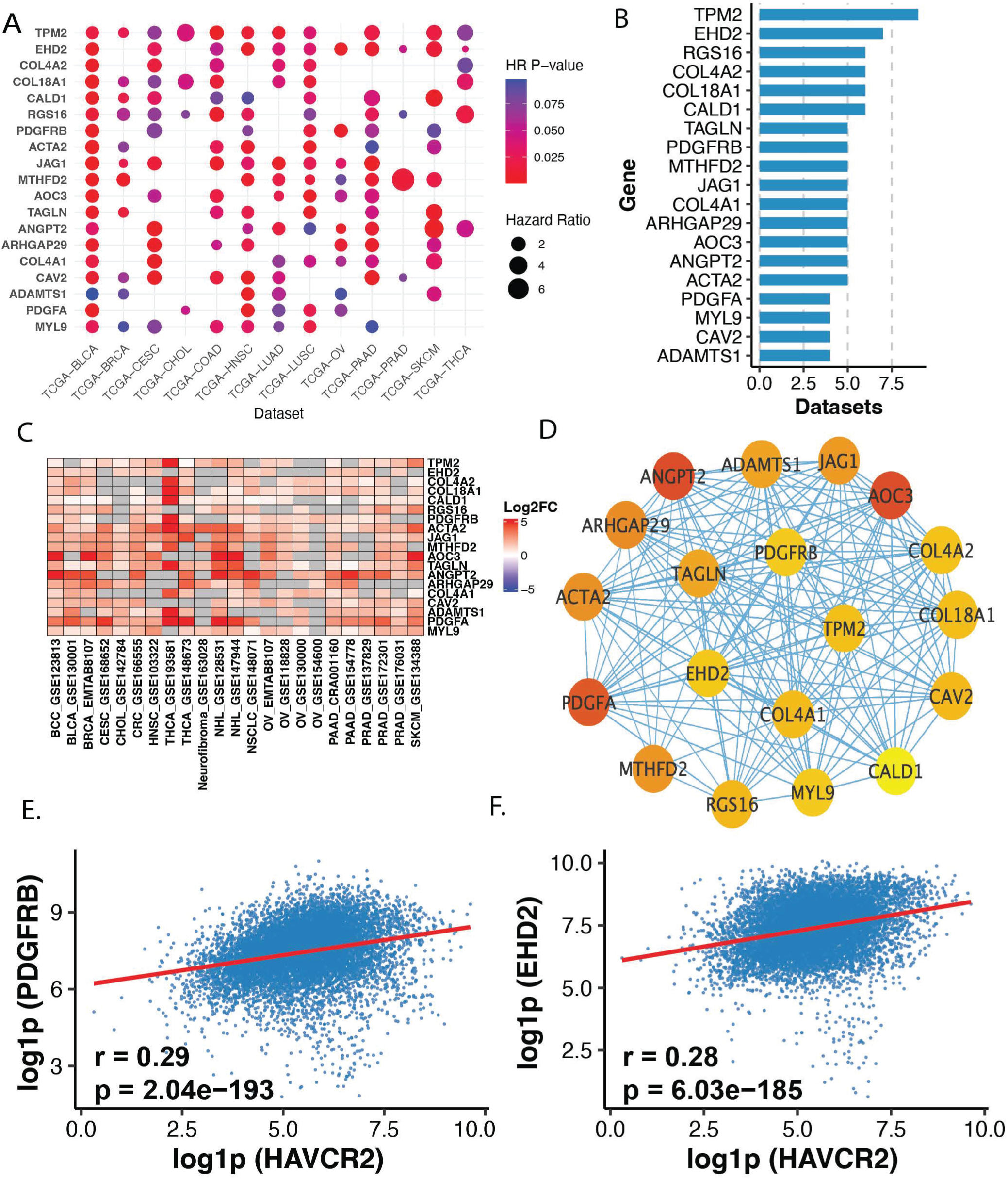
Pan-cancer pericyte gene signature associated with poor cancer outcome. **A.** Dot plot showing pan-cancer pericyte gene signature associated with poor survival outcomes in more than 30% of cancer datasets. Each row represents a cancer type from TCGA (TCGA-BLCA= Bladder Urothelial Carcinoma, TCGA-BRCA= Breast Invasive Carcinoma, TCGA-CESC= Cervical Squamous Cell Carcinoma and Endocervical Adenocarcinoma, TCGA-CHOL= Cholangiocarcinoma, TCGA-COAD= Colon Adenocarcinoma, TCGA-HNSC= Head and Neck Squamous Cell Carcinoma, TCGA-LUAD= Lung Adenocarcinoma, TCGA-LUSC= Lung Squamous Cell Carcinoma, TCGA-OV= Ovarian Serous Cystadenocarcinoma, TCGA-PAAD= Pancreatic Adenocarcinoma, TCGA-PRAD= Prostate Adenocarcinoma, TCGA-SKCM= Skin Cutaneous Melanoma, and TCGA-THCA= Thyroid Carcinoma) and each dot represents a gene. The size of the dot corresponds to the hazard ratio derived from the Cox proportional hazards model, while the colour indicates the log-rank p-value. **B.** Bar plot illustrating the frequency of poor prognosis associations for each gene across cancer types. Genes such as TPM2 and EHD2 are associated with poor outcomes in over 50% of datasets analyzed. **C.** Heatmap displaying log2 fold change of pericyte genes (associated with poor survival based on hazard ratio from Cox Hazard proportional model) across single-cell datasets of different cancers: thyroid, colon, breast, bladder, pancreatic, ovarian, prostate, head and neck squamous cell carcinoma, cholangiocarcinoma, cervical, lymphoma, lung; and one benign tumor (neurofibroma). Each row corresponds to a gene and each column to a dataset. The heatmap is color-coded by log2 fold change, with red indicating high expression. BCC= basal cell carcinoma; BLCA= bladder urothelial carcinoma; CESC= Cervical Squamous Cell Carcinoma and Endocervical Adenocarcinoma; CHOL= cholangiocarcinoma; CRC= colorectal cancer; HNSC= Head & Neck Squamous Cell Carcinoma; THCA= thyroid cancer; NHL= non-Hodgkin Lymphoma; NSCLS= non-small cell lung cancer; OV= ovary cancer; PAAD= pancreatic adeno carcinoma; PRAD= prostate adenocarcinoma; and SKCM= skin cutaneous melanoma. **D.** Gene interaction network of pan-cancer pericyte signature genes associated with poor prognosis. The network was generated using GeneMANIA, incorporating gene expression, pathway, and literature-based interactions. Each node represents a gene, and edges represent functional interactions. The network demonstrates that these pericyte genes are highly interconnected, suggesting coordinated roles in driving poor outcomes. **E–F.** Scatter plots showing representative examples of positive correlations between expression of pan-cancer pericyte signature genes and T cell exhaustion markers, such as HAVCR2. The data suggest that increased expression of pericyte genes is associated with elevated T cell exhaustion, potentially contributing to poor patient outcomes. Additional gene-level correlations are presented in Supplementary Figure 23. This analysis has been performed by downloading normalized and batch-corrected pan-cancer cancer bulk sequencing data from 10,967 samples from 32 studies (Supplementary Table 7) using cBioPortal (https://www.cbioportal.org).

To further explore their role, we assessed the correlation between the expression of these 19 genes and established markers of T cell exhaustion. Strikingly, most of the genes (13 out of 19) showed a positive correlation with exhaustion markers, suggesting that pericyte-enriched tumors may actively promote T cell exhaustion **(Fig.6E-F, Suppl. Fig. 23, Suppl. Table 7)**. This observation is consistent with our findings in **Fig.3**, where pericytes in thyroid cancer appear to interact with immune cells (CD8 T-cells) and potentially modulate their activity.

This pan-cancer pericyte signature of 19 genes provides a robust signature for precise annotation of pericytes in single-cell datasets and may serve as a foundation for future studies aimed at therapeutically targeting pericytes to improve cancer outcomes. We propose that pericytes may function as tumor shielders, similar to cancer-associated fibroblasts or cancer-associated macrophages, but remain understudied. This raises the need for comprehensive studies to map the molecular landscape of pericytes and evaluate their therapeutic potential across cancers.

### Differential gene expression of tyrosine kinases (TK) receptors across different cell types in multiple cancers

We further analyzed TK receptors gene expression across all single cell clusters because of their importance to be druggable in patients with cancer, and the clinical impact of TK inhibitors (TKI) in patients with advanced disease ^89,90^. The expression analysis revealed significantly higher expression levels of some TK receptors in pericytes compared to fibroblasts in different cancers and one benign tumor (neurofibroma) **(Fig.7)**. In particular, PDGFRβ was significantly over-expressed in the pericytes of nearly all cancers followed by NTRK2 and NTRK3 **(Fig.7)**. These findings support the notion that e.g. PDGFRβ positive pericytes in thyroid and other cancers may serve as a predictive biomarker for TKI responsiveness, offering a cellular target whose expression pattern could guide therapeutic decisions and improve patient stratification in TKI-based treatment regimens.

**Figure 7.**
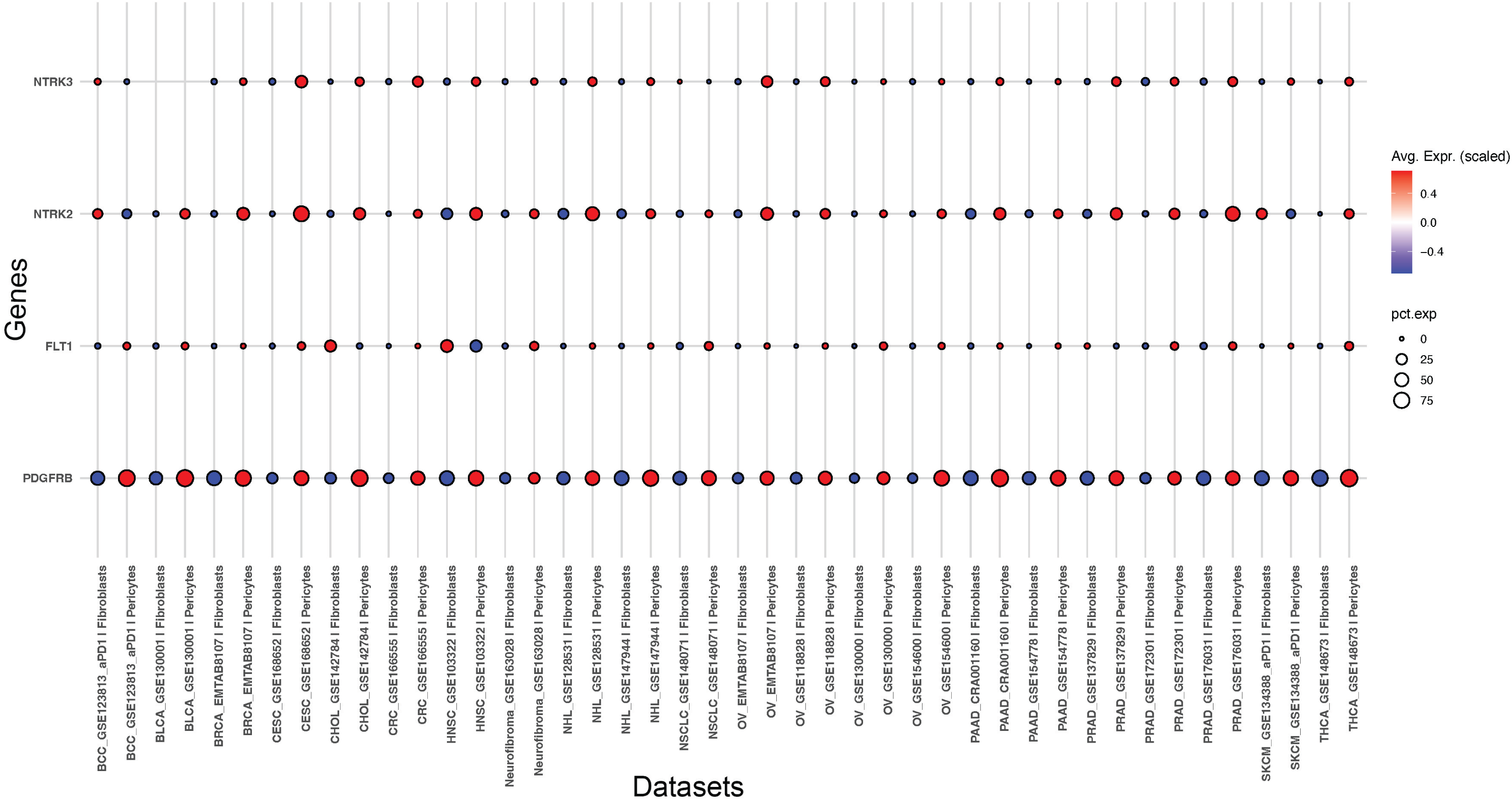
Tyrosine kinase (TK) receptors gene expression across different cancers. Dot plot illustrating the overexpression of tyrosine kinase (TK) receptors genes in pericytes compared to fibroblasts across datasets analyzed in this study. Dot color represents expression level (red = higher, blue = lower), while dot size indicates the proportion of cells expressing each gene. The TK receptors with Benjamini–Hochberg (BH) corrected p-value <0.05 and over-expression in pericytes vs fibroblasts (Log FC>0) in at least 4 datasets out of 23 were considered for this analysis. The analysis performed on following cancer: Bladder (BLCA_GSE130001), Breast (BRCA_EMTAB8107), Cervical (CESC_GSE168652), Cholangiocarcinoma (CHOL_GSE142784), Colorectal (CRC_GSE166555), Head & Neck (HNSC_GSE103322), Lymphoma (NHL_GSE128533, NHL_GSE147941), Lung (NSCLC_GSE148071), Ovarian (OV_EMTAB8107, OV_GSE118828, OV_GSE130000, OV_GSE154600), Pancreatic (PAAD_CRA001160, PAAD_GSE154778), Prostate (PRAD_GSE137629, PRAD_GSE137829, PRAD_GSE172301, PRAD_GSE176031), Skin (BCC_GSE123813_aPD1, SKCM_GSE134388_aPD1), Thyroid (THCA_GSE148673); and in one benign tumor (Neurofibroma_GSE16028).

## Discussion

Our study identifies and comprehensively characterizes pericytes in clinical samples through an extensive analysis of scRNA-seq data from different cancers. We hypothesized that pericytes play a critical role in allowing cancer cells immune escape within the tumor microenvironment, contributing to poor clinical outcomes. However, the precise mechanisms by which pericytes initiate immune escape remain poorly understood, as genome-scale studies often fail to detect and characterize these cells accurately. We have recently found for the first time that thyroid cancer (PTC and ATC) showed abundance in pericytes as compared to NT samples using bulk RNA-sequencing analyses^13,15^. Importantly, an independent study has also recently confirmed the abundance of pericyte in thyroid cancer, i.e. their enrichment in the fibrovascular core of the thyroid TME ^91^. Importantly, here in our study, we used SC-RNA-sequencing that enabled the precise detection and characterization of pericytes by directly comparing their gene expression patterns with fibroblasts, thereby generating a definitive pericyte signature. Through an analysis of publicly-available single-cell data set starting from thyroid cancers (PTC and ATC) and followed by all other cancers, we successfully manually annotated pericytes, distinguishing them from automatically annotated fibroblasts. While pericytes may be confused with fibroblasts due to phenotypic similarities, our findings reveal that these two cell types have distinct transcriptomic profiles. Single-cell and cancer transcriptome analysis empowered the derivation of pericyte gene signatures across different cancer types. By performing a comparative analysis across these datasets, we have identified a robust pan-cancer pericyte-associated gene signature. This signature includes well-known pericyte markers such as ACTA2, and also uncovers multiple additional genes that are not widely recognized in the context of pericyte biology. These newly identified genes may represent novel components of the pericyte transcriptional program within the TME, offering opportunities for deeper mechanistic insights and potential therapeutic targeting.

The single-cell analysis of cell communication revealed, for the first time ever, that pericytes act as central signaling hubs in the tumor ecosystem of thyroid cancer. These cells engage extensively with immune cells (e.g., T cells) and cancer cells, and to a lesser extent with stromal and vascular cells (e.g., fibroblasts, endothelial cells), suggesting that pericytes act as master regulators in shaping the tumor ecosystem to promote cancer growth. E.g. the COL4A1 on pericytes interacts with the CD44 receptor on T cells; CD44 is known to play a key role in T cell activation and infiltration to eradicate tumors ^49,92^. Additionally, Nectin-2 over-expressed from pericytes can bind to TIGIT on a T cell, preventing the T cell from effectively attacking tumor cells. This overexpression and interaction of Nectin-2 with TIGIT have been identified across many other cancers, including breast and ovarian cancers to suppress the immune response. Nectin-2 binding to TIGIT inhibits T cell proliferation and cytokine production, impairing NK cell cytotoxicity and resulting in tumour growth and progression ^93–95^. Targeting the Nectin-2 and TIGIT-axis is a promising therapeutic strategy, with multiple clinical trials underway to assess the efficacy of anti-TIGIT antibodies in restoring antitumor immunity ^93–95^. Further supported survival analysis on pericytes genes demonstrated that the enrichment of pericytes correlated with poor clinical outcomes in thyroid cancer. These last results also align with similar findings in other cancers, including colon, head & neck, bladder, pancreatic, lung, breast, ovarian, cervical, prostate and melanoma where pericyte abundance is associated with aggressive disease and unfavorable prognosis. Collectively, these results suggest that pericytes may promote tumor progression by facilitating immune evasion, thereby shielding cancer cells from immune surveillance and enabling tumor growth.

The findings presented here indicate that targeting pericytes within the TME could disrupt the immunosuppressive niche, potentially enhancing T cell-mediated anti-tumor responses. By identifying the gene expression signatures of pericytes and their communication pathways, this study provides a valuable resource for developing novel therapeutic strategies and biomarkers aimed at targeting pericytes. Additionally, the enrichment of pericytes in cancers might help in improved patient stratification, enabling the identification of high-risk individuals who may benefit from more intensive treatment and follow-up.

## Conclusions

This large-scale scRNA-seq analysis of multiple datasets defines a pan-cancer pericyte signature and sheds light on pericyte activity, particularly their communication with T cells, suggesting a role in T cell exhaustion, a key factor in the failure of many cancer therapies.

This study underscores the pivotal role of pericyte population enhancement as a critical regulator of the TME, facilitating immune evasion, and driving disease progression. TK expression in pericytes may serve as a predictive biomarker for TKI responsiveness, offering a cellular target whose expression pattern may guide therapeutic decisions and improve patient stratification in TKI-based treatment regimens.

### Translational significance

These findings provide clinically actionable insights by identifying pericyte-derived molecular targets that may refine patient risk stratification and support personalized treatment approaches. The pericyte-associated markers uncovered here show strong prognostic value across multiple cancer types, highlighting their potential to improve outcome prediction. Moreover, the specificity of these targets positions them as promising candidates for therapeutic intervention and for incorporation into future biomarker-driven clinical trials.

## Acknowledgments

We thank those authors whom we were not able to cite because of limited space. We gratefully acknowledge Dr. Andrea Calabria, Director of Computational Biology (Tessera Therapeutics), for his critical reading of the manuscript. We also thank Dr. Alessandro Prete (BIDMC) for his critical reading.

## Authors’ Contributions

Conceptualization: CN

Experimental design: AB, CN

Writing the manuscript: AB, CN

Substantive editing of the manuscript: AB, YB, JL, CN

Literature review: AB, YB, JL, CN

Computational analyses: AB

Supervision: CN

## Note

This research meets the ethics guidelines. Data sharing is not applicable to this article as datasets were already publicly-available from other data sets (Suppl. Table 3). SC-RNA-seq data were downloaded from different datasets from Gene Expression Omnibus (GEO) or Tumor Immune single-cell Hub.

