## Supplementary material for "Single-cell analysis reveals a universal pericyte signature associated with poor clinical outcome and immune T cell dysfunction in thyroid cancer and other cancers": Suppl. Figures

**Supplementary Figures:**

**A.**

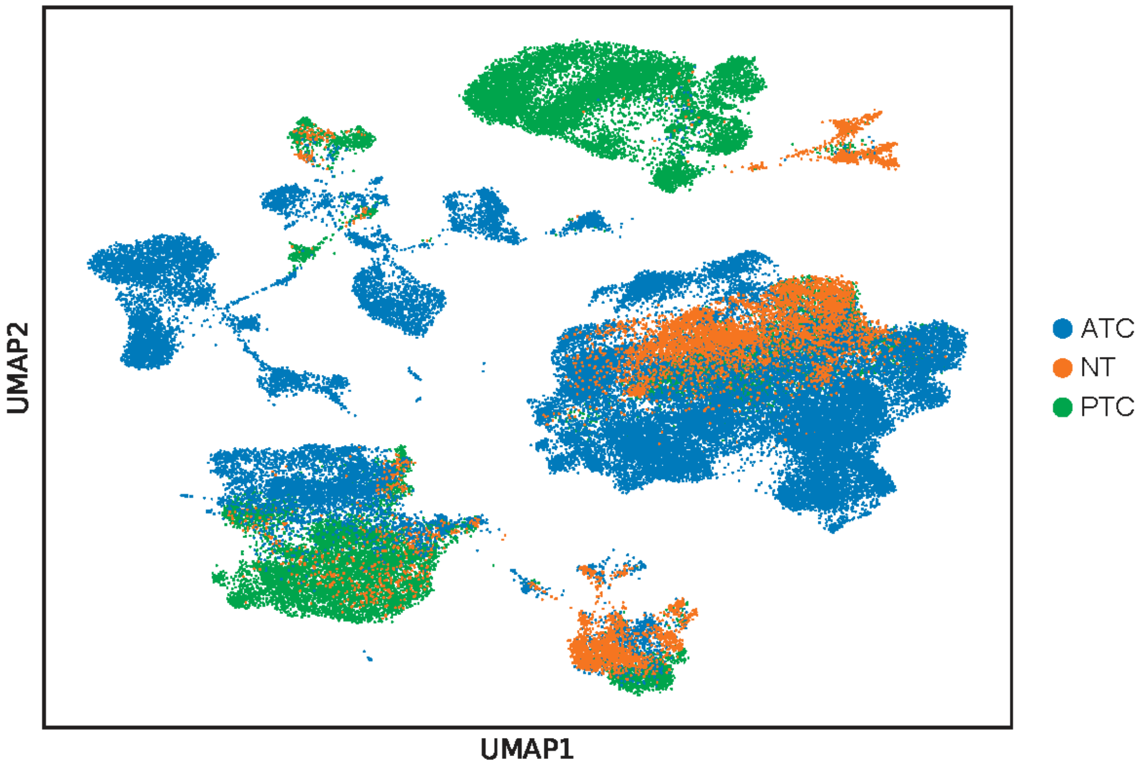

B.

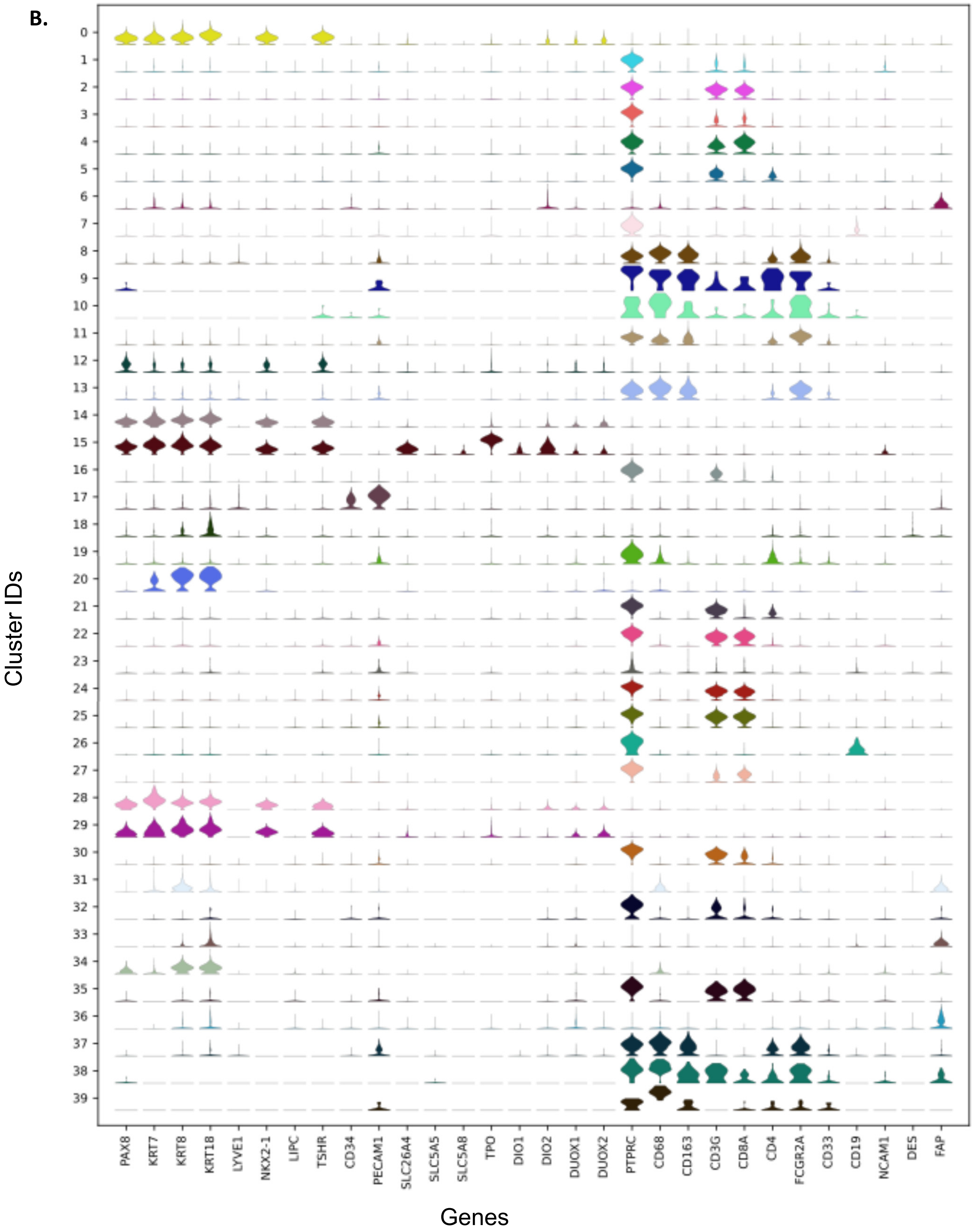
Cluster IDs
PAX8
KRT7
KRT8
KRT18
LYVE1
NKX2-1
LIPC
TSHR
CD34
PECAM1
SLC26A4
SLC5A5
SLC5A8
TPO
DIO1
DIO2
DUOX1
DUOX2
PTPRC
CD68
CD163
CD3G
CD8A
CD4
FCGR2A
CD33
CD19
NCAM1
DES
FAP
Genes

**Supplementary Fig. 1. Analysis of thyroid cancer dataset (GSE193581): A.** UMAP of single-cell transcriptomic data from the thyroid cancer dataset, colored by clinical sample group. ATC represents anaplastic thyroid cancer, PTC denotes papillary thyroid cancer, and NT corresponds to non-cancerous thyroid tissue samples. **B**. Violin plots showing the normalised raw gene expression of thyroid, immune, and stromal genes in each cluster manually annotated (0 = PTC_Tumor, 1 = T-Cells:CTL, 2 = T-Cells, 3 = T-Cells, 4 = T-Cells:CTL, 5 = T-Cells, 6 = ATC_Tumor, 7 = B-Cells, 8 = Monocytes_Macro, 9 = Monocytes_Macro, 10 = Monocytes_Macro, 11 = Monocytes_Macro, 12 = PTC_Tumor, 13 = Monocytes_Macro, 14 = PTC_Tumor, 15 = PTC_Tumor, 16 = T-Cells, 17 = Endothelial, 18 = Fibroblasts, 19 = DB, 20 = PTC_Tumor, 21 = T-Cells, 22 = T-Cells:CTL, 23 = B-Cells, 24 = T-Cells:CTL, 25 = T-Cells:CTL, 26 = B-Cells, 27 = T-Cells:CTL, 28 = PTC_Tumor, 29 = PTC_Tumor, 30 = T-Cells:CTL, 31 = ATC_Tumor, 32 = Fibroblasts, 33 = ATC_Tumor1, 34 = ATC_Tumor, 35 = T-Cells:CTL, 36 = ATC_Tumor1, 37 = Fibroblasts, 38 = DB, 39 = Monocytes_Macro).

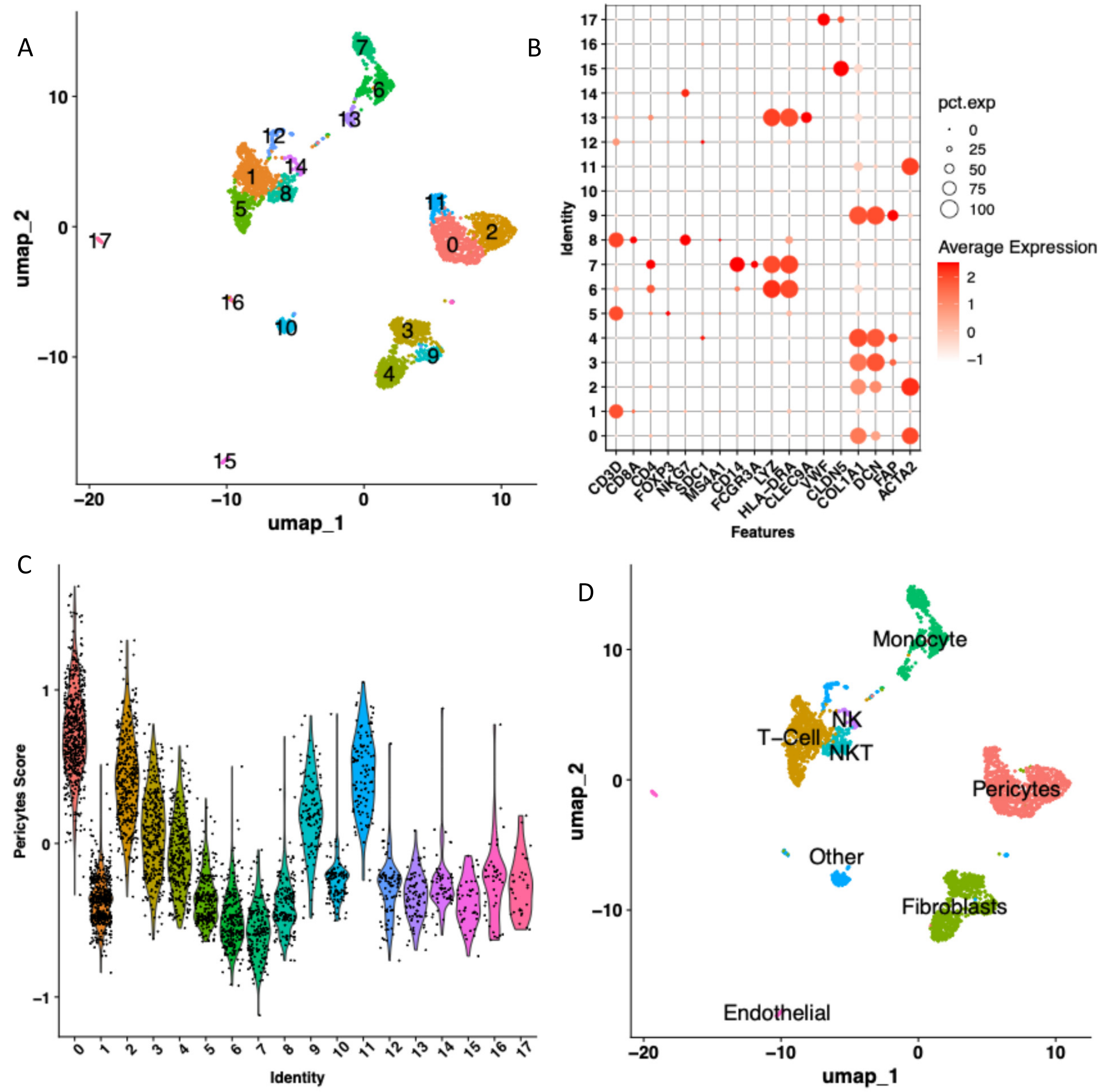

**Supplementary Figure 2. Analysis of melanoma Dataset (GSE134388): A.** UMAP projection of single-cell transcriptomic data, with cells colored by unsupervised cluster IDs. **B.** Dot plot showing expression of canonical cell type markers across clusters. **C.** Expression of pericyte-related markers across clusters based on module score calculation using genes from Table 1 and the Seurat Package. Clusters IDs (0, 2, 11) have high pericytes signature score therefore annotated as pericytes. 0=Pericytes, 1=T-Cell, 2=Pericytes, 3=Fibroblasts, 4=Fibroblasts, 5=T-Cell, 6=Monocyte, 7=Monocyte, 8=NKT, 9=Fibroblasts, 10=Other, 11=Pericytes, 12=Other, 13=Monocyte, 14=NK, 15=Endothelial, 16=Other, 17=Endothelial. **D.** Cell type annotations based on canonical marker expression and pericyte gene signature.

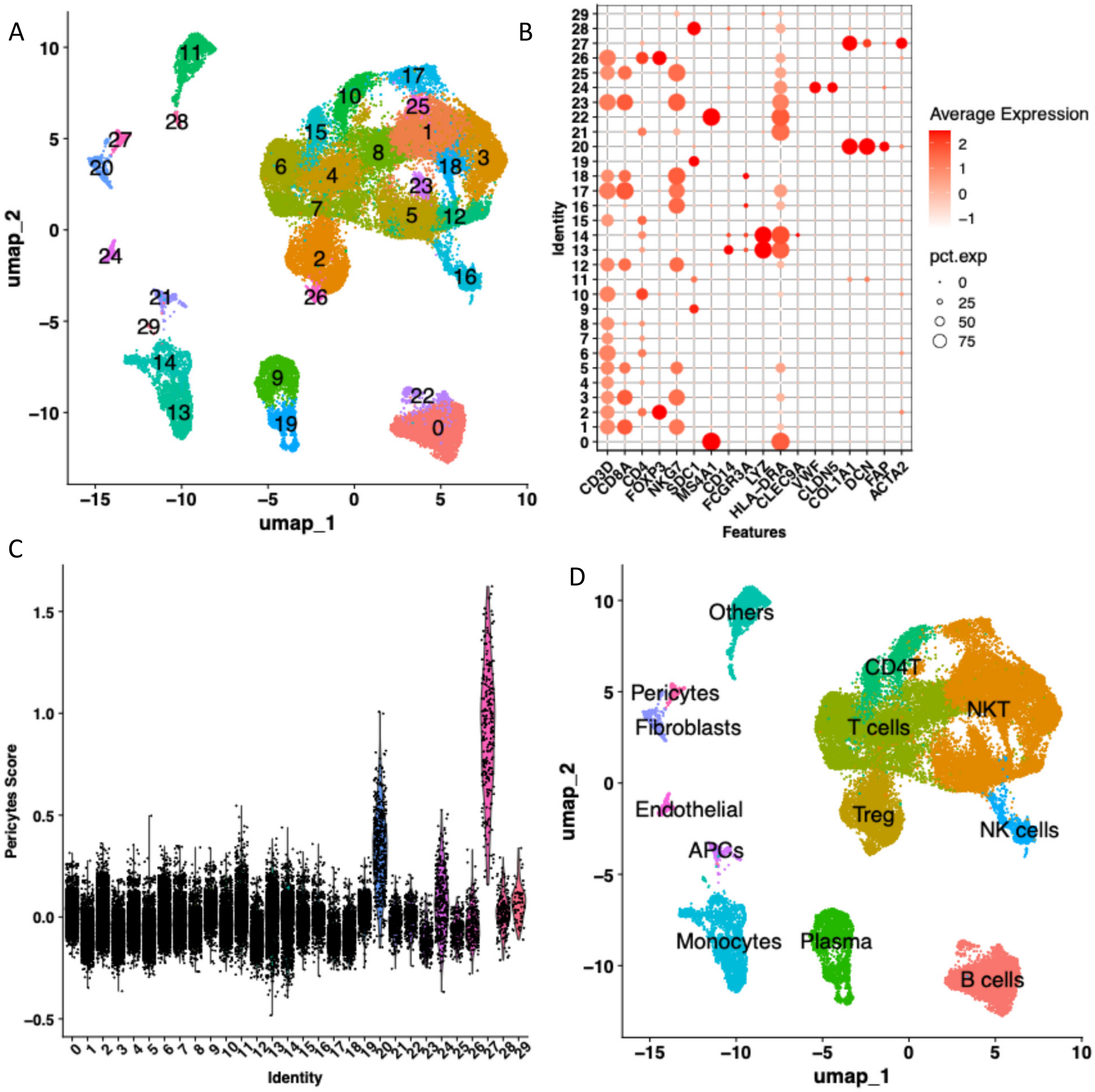

**Supplementary Figure 3. Analysis of melanoma dataset (GSE 123813): A.** UMAP projection of single-cell transcriptomic data, with cells colored by unsupervised cluster IDs. **B.** Dot plot showing expression of canonical cell type markers across clusters. **C.** Expression of pericyte-related markers across clusters based on module score calculation using genes from Table 1 and the Seurat Package. Cluster ID #27 has high pericytes signature score therefore annotated as pericytes. 0=B cells, 1=NKT, 2=Treg, 3=NKT, 4=T cells, 5=NKT, 6=T cells, 7=T cells, 8=T cells, 9=Plasma, 10=CD4T, 11=Others, 12=NKT, 13=Monocytes, 14=Monocytes, 15=CD4T, 16=NK cells, 17=NKT, 18=NKT, 19=Plasma, 20=Fibroblasts, 21=APCs, 22=B cells, 23=NKT, 24=Endothelial, 25=NKT, 26=Treg, 27=Pericytes, 28=Others, 29=Others. **D.** Cell type annotations based on canonical marker expression and pericyte gene signature.

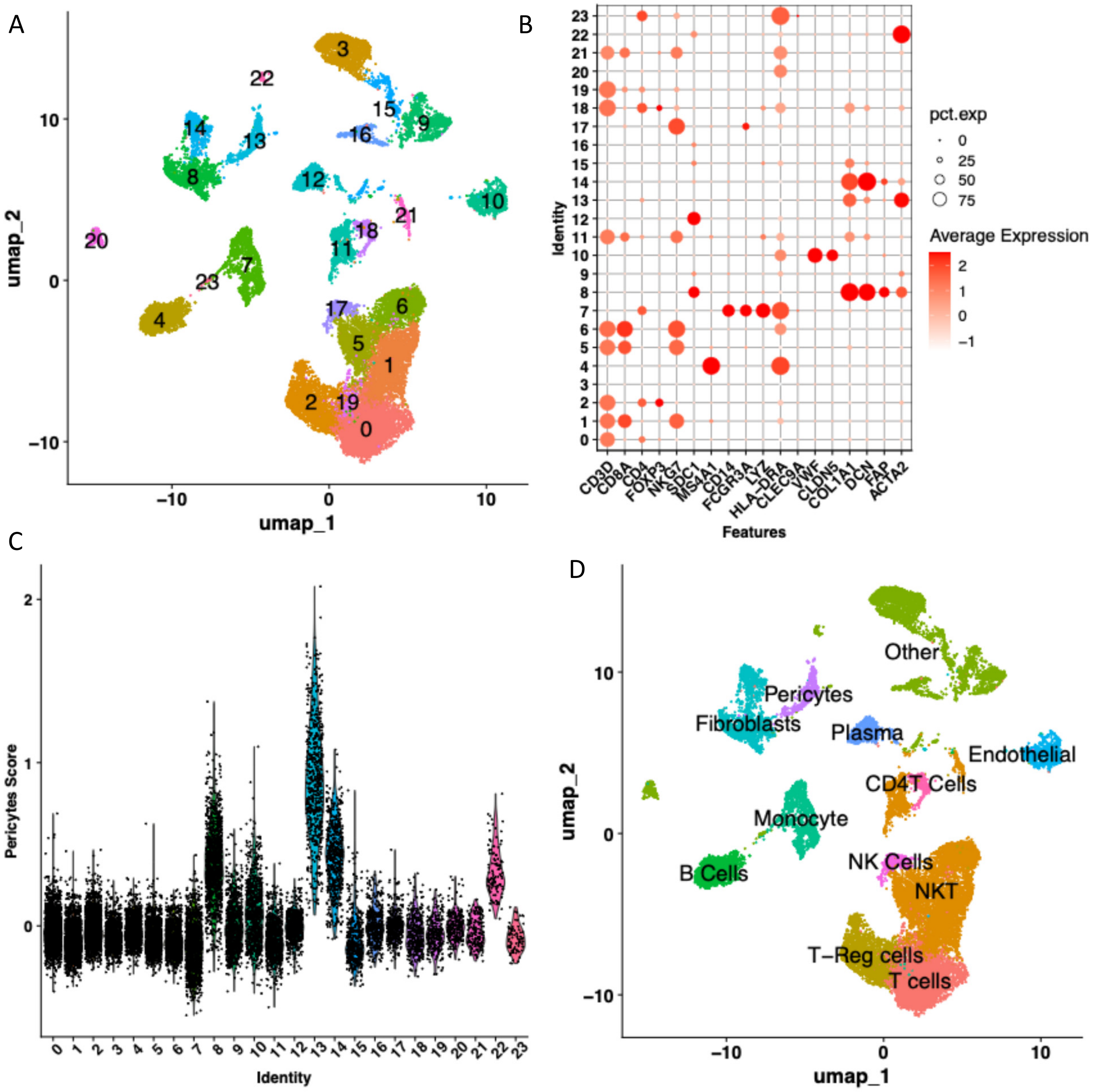

**Supplementary Figure 4. Analysis of breast cancer dataset (EMTAB8107): A.** UMAP projection of single-cell transcriptomic data, with cells colored by unsupervised cluster IDs. **B.** Dot plot showing expression of canonical cell type markers across clusters. **C.** Expression of pericyte-related markers across clusters based on module score calculation using genes from Table 1 and the Seurat Package. Cluster ID #13 has high pericytes signature score therefore annotated as pericytes. =T cells, 1=NKT, 2=T-Reg cells, 3=Other, 4=B Cells, 5=NKT, 6=NKT, 7=Monocyte, 8=Fibroblasts, 9=Other, 10=Endothelial, 11=NKT, 12=Plasma, 13=Pericytes, 14=Fibroblasts, 15=Other, 16=Other, 17=NK Cells, 18=CD4T Cells, 19=T cells, 20=Other, 21=NKT, 22=Other, 23=Other. **D.** Cell type annotations based on canonical marker expression and pericyte gene signature.

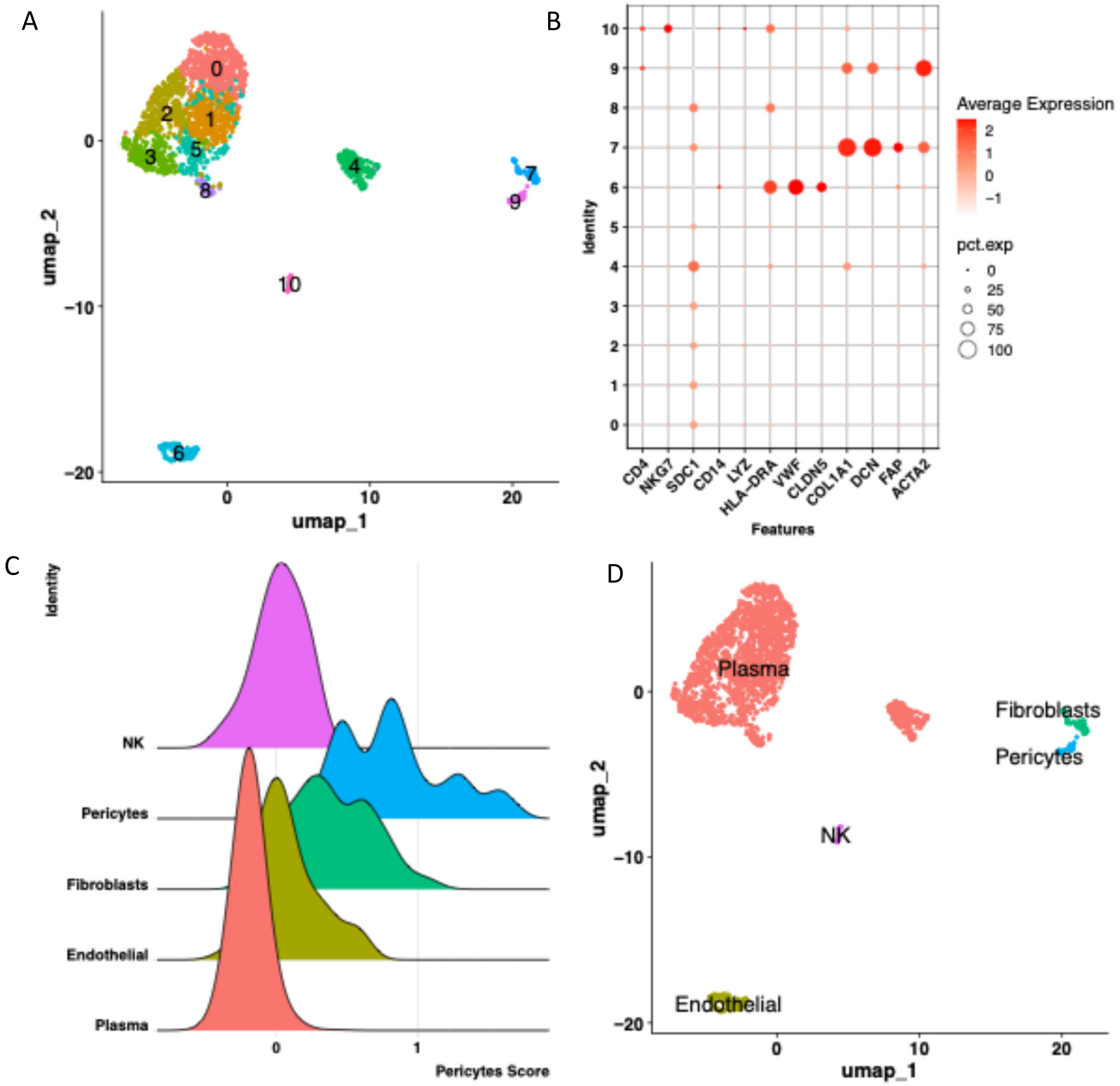

**Supplementary Figure 5. Analysis of bladder cancer dataset (GSE130001): A.** UMAP projection of single-cell transcriptomic data, with cells colored by unsupervised cluster IDs. **B.** Dot plot showing expression of canonical cell type markers across clusters. **C.** Expression of pericyte-related markers across clusters based on module score calculation using genes from Table 1 and the Seurat Package. Cluster ID #9 has high pericytes signature score therefore annotated as pericytes. =Plasma, 1=Plasma, 2=Plasma, 3=Plasma, 4=Plasma, 5=Plasma, 6=Endothelial, 7=Fibroblasts, 8=Plasma, 9=Pericytes, 10=NK, **D.** Cell type annotations based on canonical marker expression and pericyte gene signature.

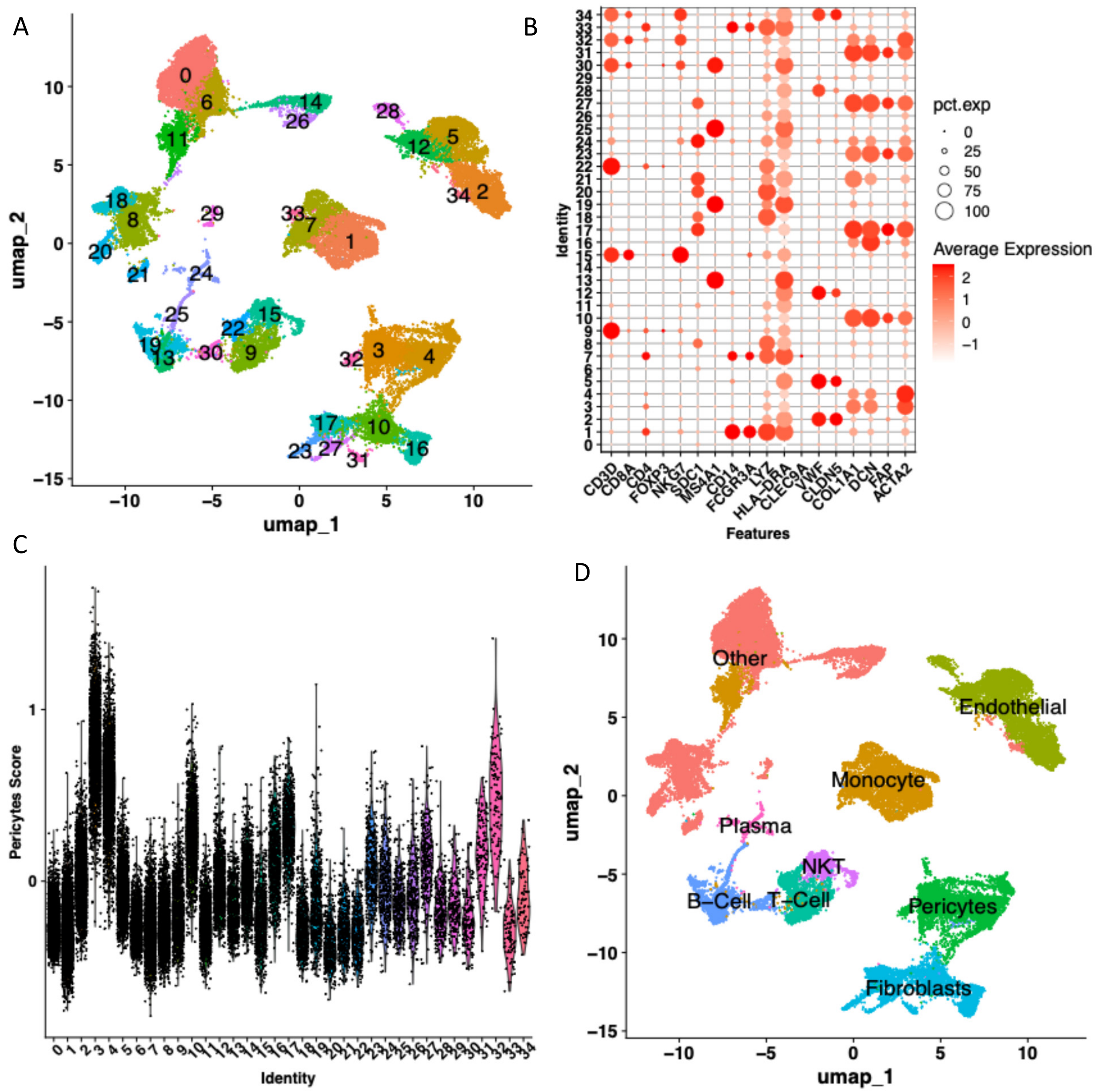

**Supplementary Figure 6. Analysis of pancreatic cancer dataset (CRA001160): A.** UMAP projection of single-cell transcriptomic data, with cells colored by unsupervised cluster IDs. **B.** Dot plot showing expression of canonical cell type markers across clusters. **C.** Expression of pericyte-related markers across clusters based on module score calculation using genes from Table 1 and the Seurat Package. Clusters with IDs #3, #4, #32 have high pericytes signature score therefore annotated as pericytes. 0=Other, 1=Monocyte, 2=Endothelial, 3=Pericytes, 4=Pericytes, 5=Endothelial, 6=Other, 7=Monocyte, 8=Other, 9=T-Cell, 10=Fibroblasts, 11=Monocyte, 12=Endothelial, 13=B-Cell, 14=Other, 15=NKT, 16=Fibroblasts, 17=Fibroblasts, 18=Other, 19=B-Cell, 20=Other, 21=Other, 22=T-Cell, 23=Fibroblasts, 24=Plasma, 25=B-Cell, 26=Other, 27=Fibroblasts, 28=Endothelial, 29=Other, 30=B-Cell, 31=Fibroblasts, 32=Pericytes, 33=Monocyte, 34=Other. **D.** Cell type annotations based on canonical marker expression and pericyte gene signature.

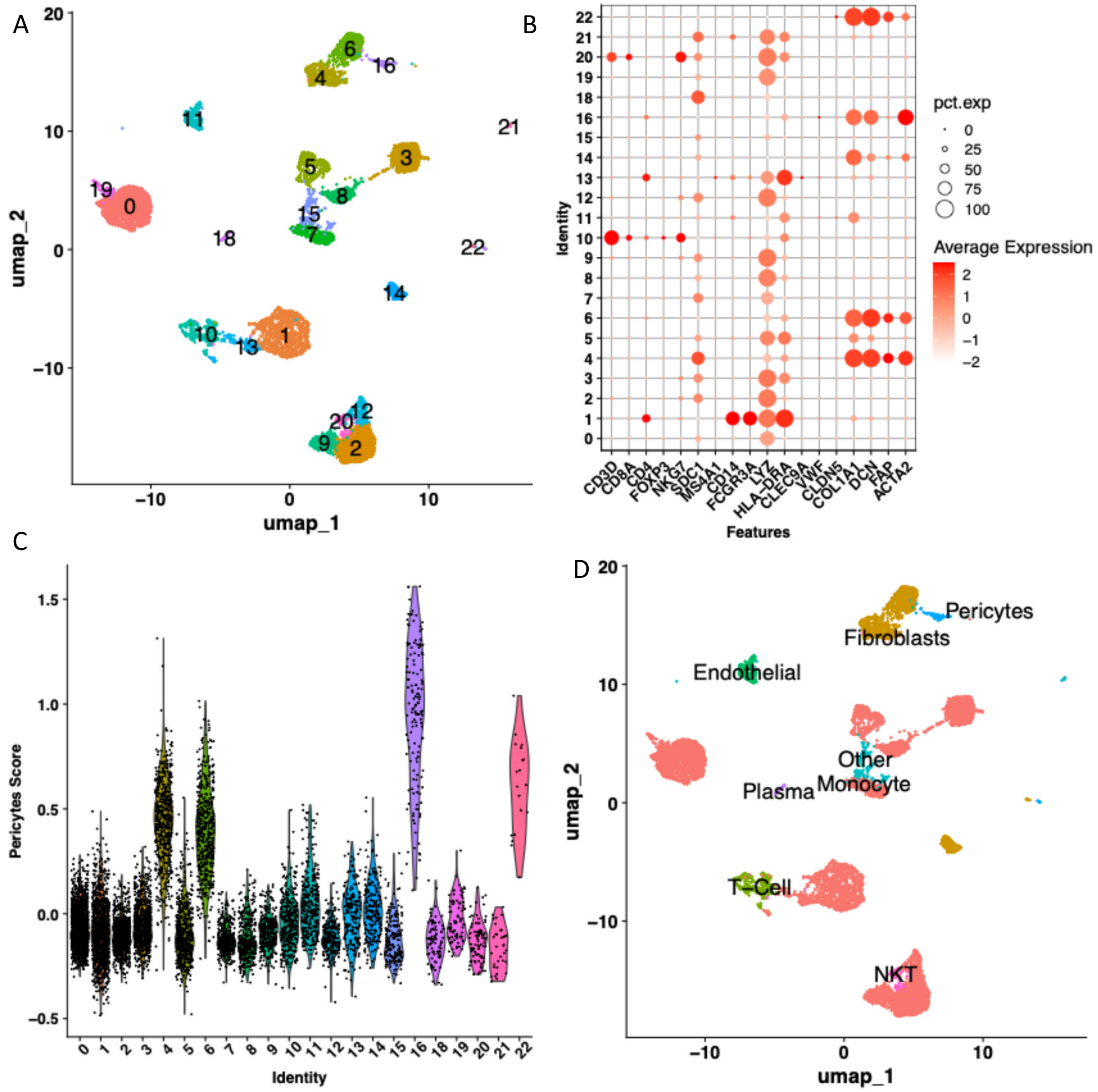

**Supplementary Figure 7. Analysis of pancreatic cancer dataset (GSE154778): A.** UMAP projection of single-cell transcriptomic data, with cells colored by unsupervised cluster IDs. **B.** Dot plot showing expression of canonical cell type markers across clusters. **C.** Expression of pericyte-related markers across clusters based on module score calculation using genes from Table 1 and the Seurat Package. Cluster ID #16 has high pericytes signature score therefore annotated as pericytes. 0=Monocyte, 1=Monocyte, 2=Monocyte, 3=Monocyte, 4=Fibroblasts, 5=Monocyte, 6=Fibroblasts, 7=Monocyte, 8=Monocyte, 9=Monocyte, 10=T-Cell, 11=Endothelial, 12=Monocyte, 13=Monocyte, 14=Fibroblasts, 15=Other, 16=Pericytes, 18=Plasma, 19=Monocyte, 20=NKT, 21=Other, 22=Fibroblasts. **D.** Cell type annotations based on canonical marker expression and pericyte gene signature.

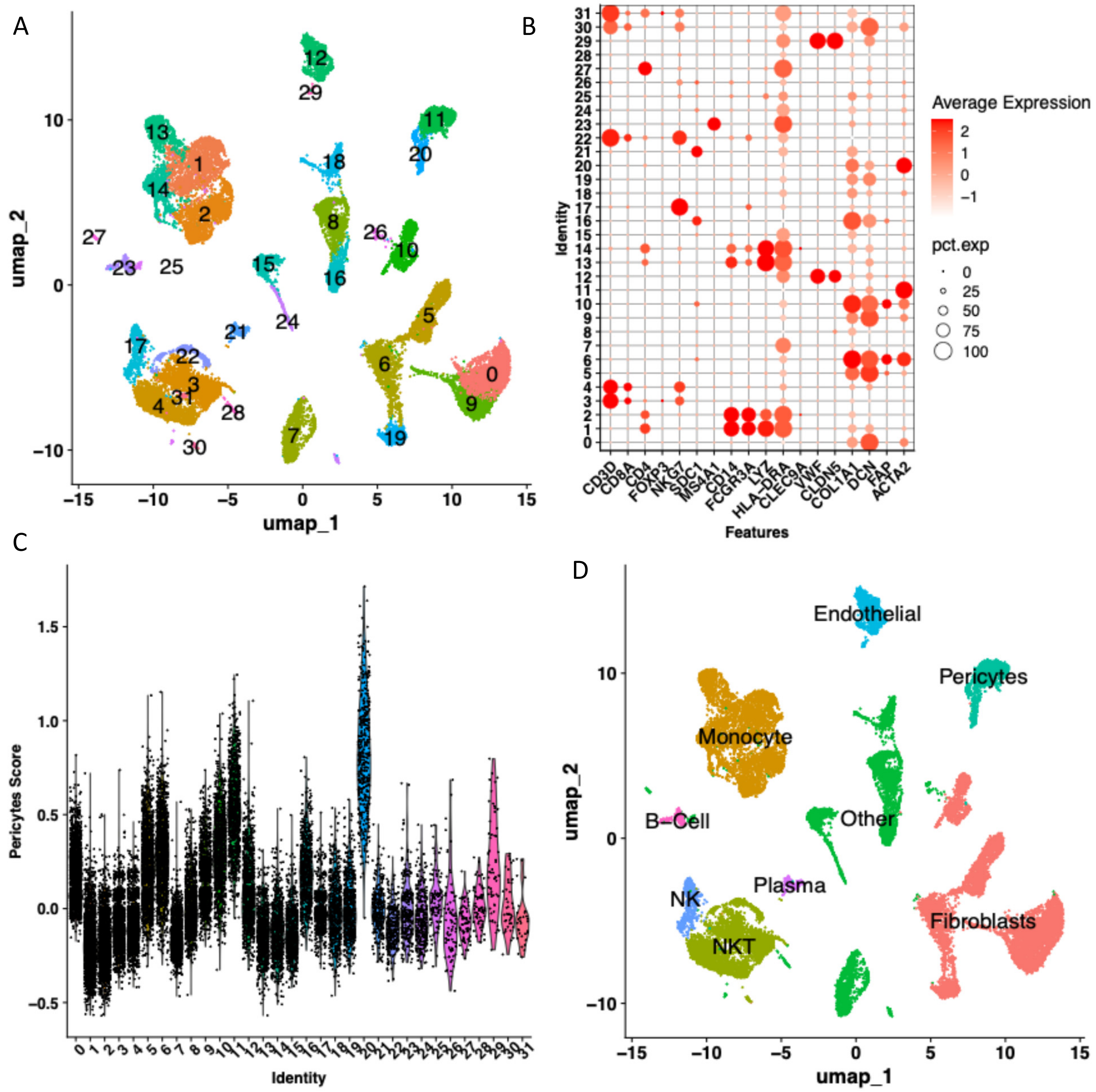

**Supplementary Figure 8. Analysis of ovarian cancer dataset (EMTAB8107): A.** UMAP projection of single-cell transcriptomic data, with cells colored by unsupervised cluster IDs. **B.** Dot plot showing expression of canonical cell type markers across clusters. **C.** Expression of pericyte-related markers across clusters based on module score calculation using genes from Table 1 and the Seurat Package. Cluster IDs #11, #20 have high pericytes signature score therefore annotated as pericytes. 0=Fibroblasts, 1=Monocyte, 2=Monocyte, 3=NKT, 4=NKT, 5=Fibroblasts, 6=Fibroblasts, 7=Other, 8=Other, 9=Fibroblasts, 10=Fibroblasts, 11=Pericytes, 12=Endothelial, 13=Monocyte, 14=Monocyte, 15=Other, 16=Other, 17=NK , 18=Other, 19=Fibroblasts, 20=Pericytes, 21=Plasma, 22=NKT, 23=B-Cell, 24=Other, 25=Other, 26=Other, 27=Other, 28=Other, 29=Endothelial, 30=NKT, 31=NKT. **D.** Cell type annotations based on canonical marker expression and pericyte gene signature.

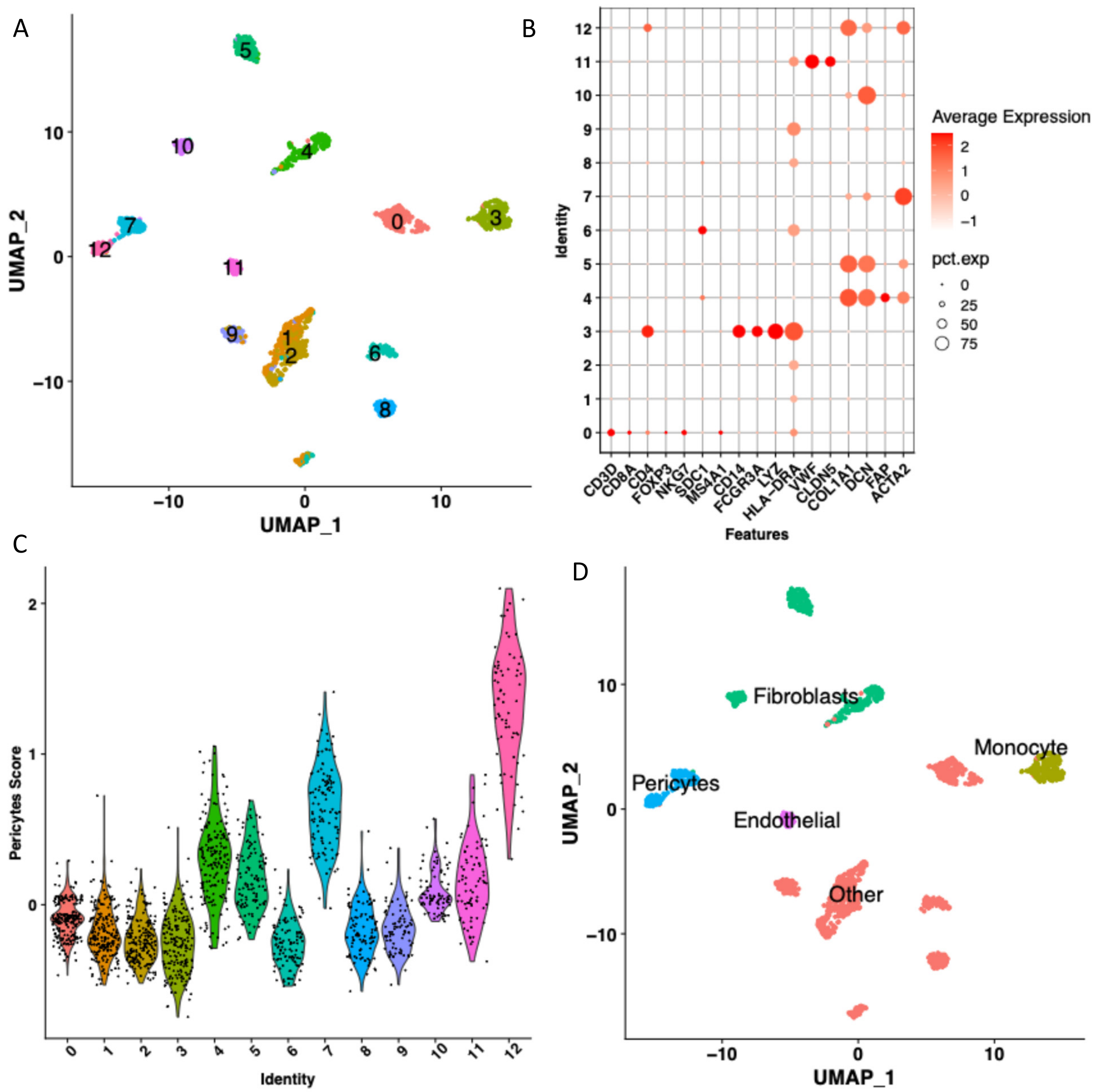

**Supplementary Figure 9. Analysis of ovarian cancer dataset (GSE118828): A.** UMAP projection of single-cell transcriptomic data, with cells colored by unsupervised cluster IDs. **B.** Dot plot showing expression of canonical cell type markers across clusters. **C.** Expression of pericyte-related markers across clusters based on module score calculation using genes from Table 1 and the Seurat Package. Cluster IDs #7, #12 have high pericytes signature score therefore annotated as pericytes. 0=Other, 1=Other, 2=Other, 3=Monocyte, 4=Fibroblasts, 5=Fibroblasts, 6=Other, 7=Pericytes, 8=Other, 9=Other, 10=Fibroblasts, 11=Endothelial, 12=Pericytes. **D.** Cell type annotations based on canonical marker expression and pericyte gene signature.

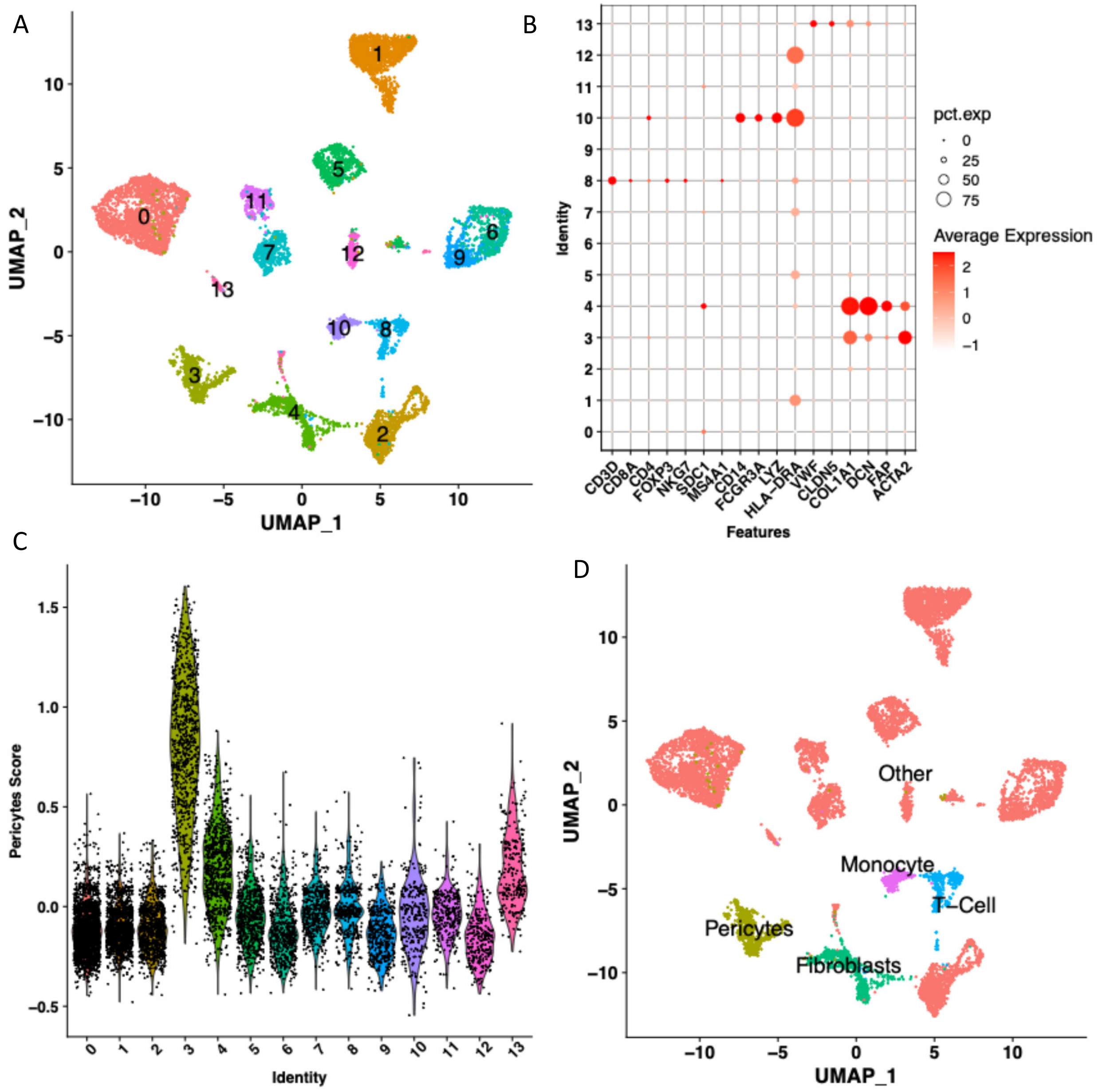

**Supplementary Figure 10. Analysis of ovarian cancer dataset (GSE130000): A.** UMAP projection of single-cell transcriptomic data, with cells colored by unsupervised cluster IDs. **B.** Dot plot showing expression of canonical cell type markers across clusters. **C.** Expression of pericyte-related markers across clusters based on module score calculation using genes from Table 1 and the Seurat Package. Cluster ID #3 has high pericytes signature score therefore annotated as pericytes. 0=Other, 1=Other, 2=Other, 3=Pericytes, 4=Fibroblasts, 5=Other, 6=Other, 7=Other, 8=T-Cell, 9=Other, 10=Monocyte, 11=Other, 12=Other, 13=Other. **D.** Cell type annotations based on canonical marker expression and pericyte gene signature.

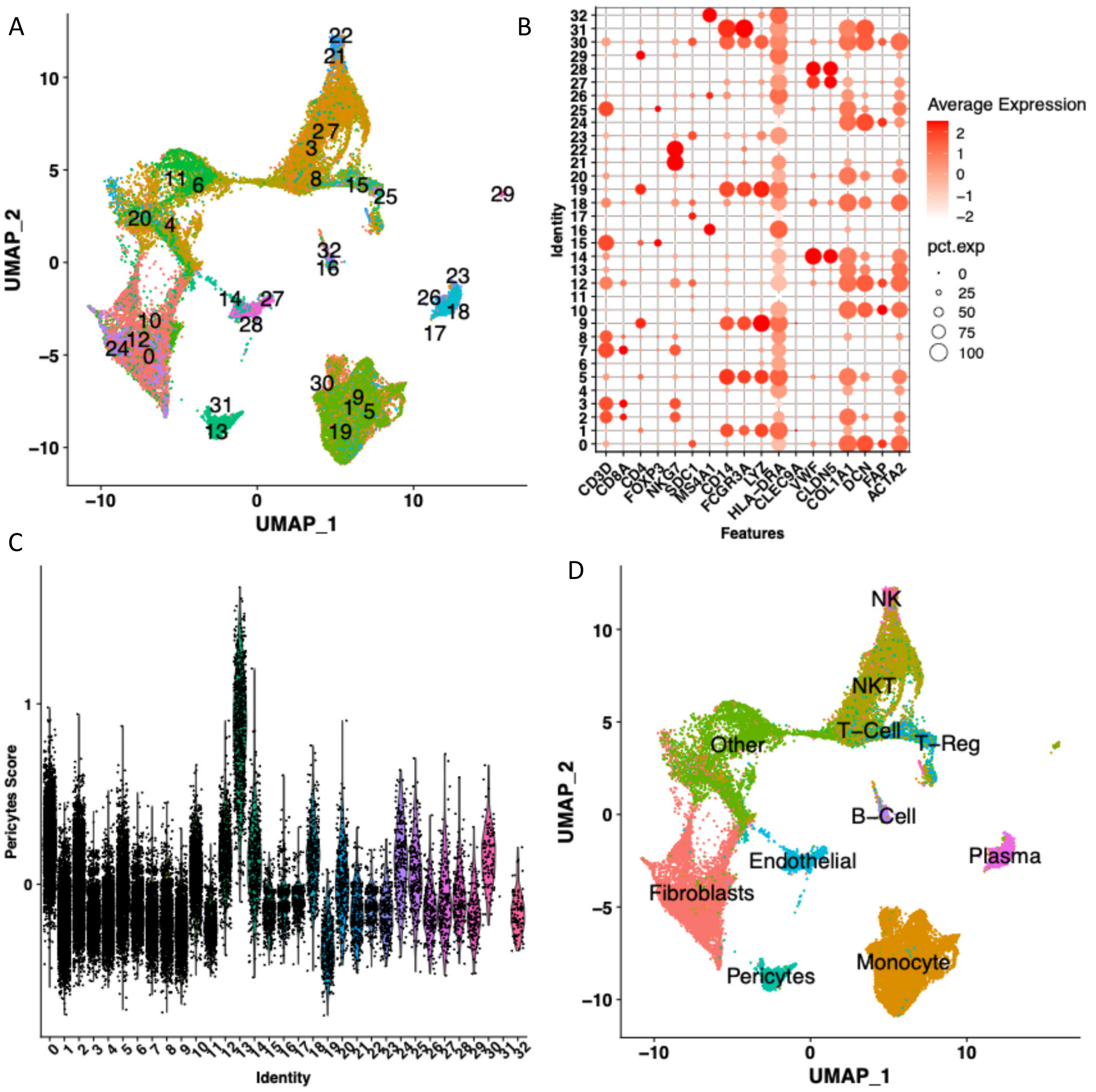

**Supplementary Figure 11. Analysis of ovarian cancer dataset (GSE154600): A.** UMAP projection of single-cell transcriptomic data, with cells colored by unsupervised cluster IDs. **B.** Dot plot showing expression of canonical cell type markers across clusters. **C.** Expression of pericyte-related markers across clusters based on module score calculation using genes from Table 1 and the Seurat Package. Cluster ID #13 has high pericytes signature score therefore annotated as pericytes. 0=Fibroblasts, 1=Monocyte, 2=NKT, 3=NKT, 4=Other, 5=Monocyte, 6=Other, 7=NKT, 8=T-Cell, 9=Monocyte, 10=Fibroblasts, 11=Other, 12=Fibroblasts, 13=Pericytes, 14=Endothelial, 15=T-Reg, 16=B-Cell, 17=Plasma, 18=Fibroblasts, 19=Monocyte, 20=Fibroblasts, 21=NK , 22=NK , 23=Plasma, 24=Fibroblasts, 25=T-Reg, 26=B-Cell, 27=Endothelial, 28=Endothelial, 29=Other, 30=Monocyte, 31=Other, 32=B-Cell. **D.** Cell type annotations based on canonical marker expression and pericyte gene signature.

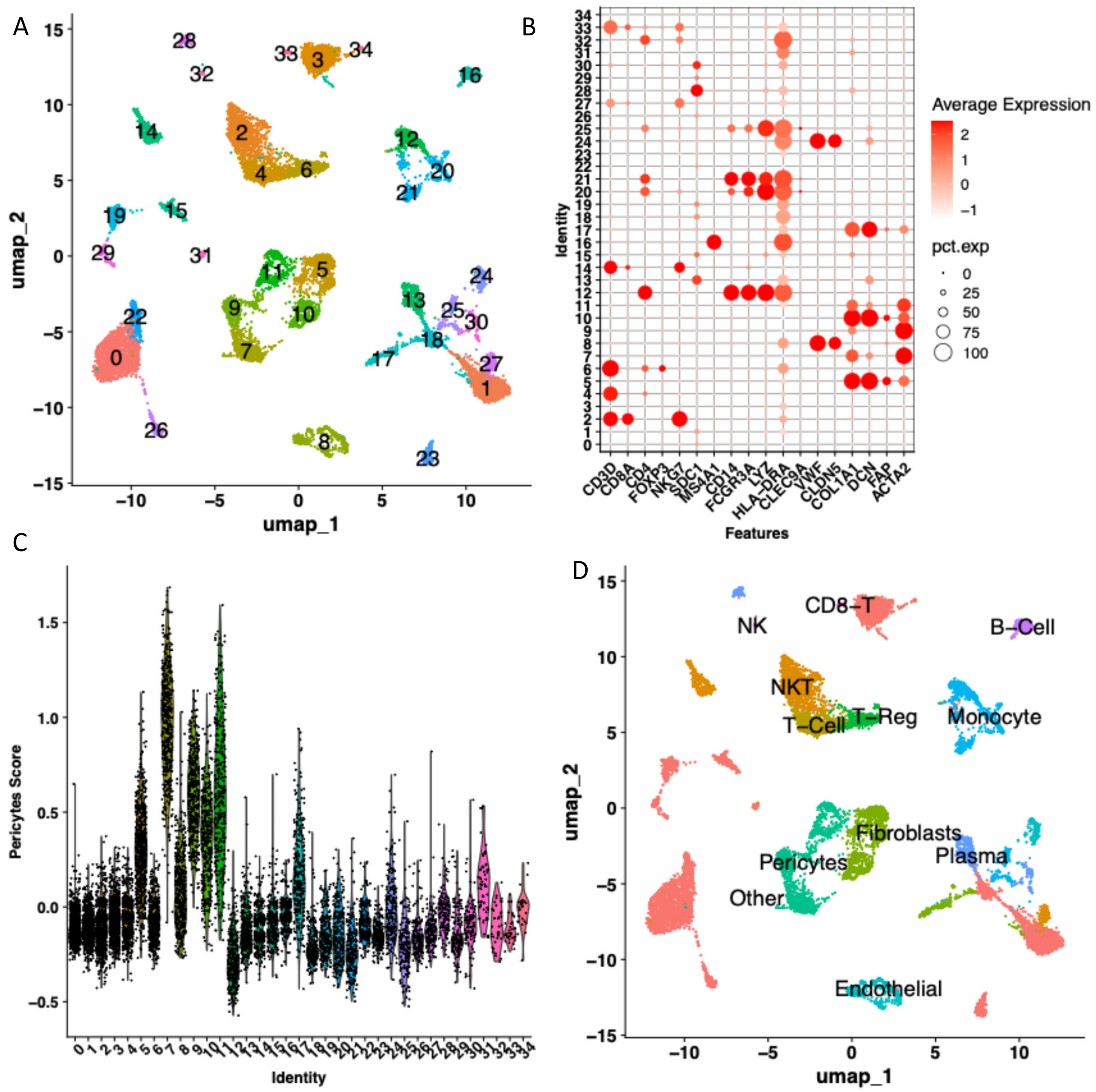

**Supplementary Figure 12. Analysis of prostate cancer dataset (GSE137829): A.** UMAP projection of single-cell transcriptomic data, with cells colored by unsupervised cluster IDs. **B.** Dot plot showing expression of canonical cell type markers across clusters. **C.** Expression of pericyte-related markers across clusters based on module score calculation using genes from Table 1 and the Seurat Package. Cluster IDs #7, #9, #11 have high pericytes signature score therefore annotated as pericytes. 0=Other, 1=Other, 2=NKT, 3=Other, 4=T-Cell, 5=Fibroblasts, 6=T-Reg, 7=Pericytes, 8=Endothelial, 9=Pericytes, 10=Fibroblasts, 11=Pericytes, 12=Monocyte, 13=Plasma, 14=NKT, 15=Other, 16=B-Cell, 17=Fibroblasts, 18=Other, 19=Other, 20=Monocyte, 21=Monocyte, 22=Other, 23=Other, 24=Endothelial, 25=Monocyte, 26=Other, 27=NKT, 28=Plasma, 29=Other, 30=Plasma, 31=Other, 32=NK , 33=CD8-T, 34=Other. **D.** Cell type annotations based on canonical marker expression and pericyte gene signature.

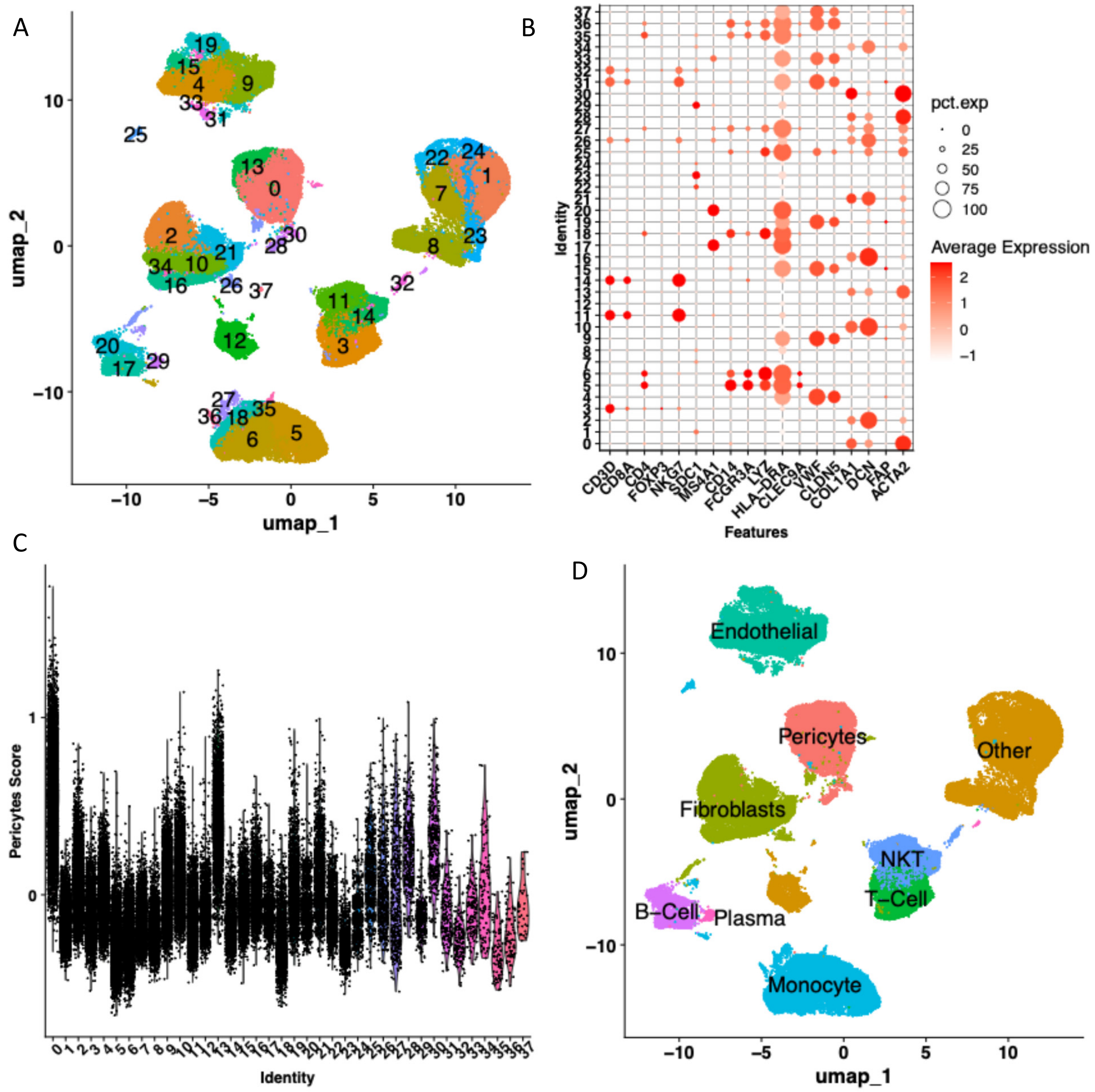

**Supplementary Figure 13. Analysis of prostate cancer dataset (GSE172301): A.** UMAP projection of single-cell transcriptomic data, with cells colored by unsupervised cluster IDs. **B.** Dot plot showing expression of canonical cell type markers across clusters. **C.** Expression of pericyte-related markers across clusters based on module score calculation using genes from Table 1 and the Seurat Package. Cluster IDs #0, #13 have high pericytes signature score therefore annotated as pericytes. 0=Pericytes, 1=Other, 2=Fibroblasts, 3=T-Cell, 4=Endothelial, 5=Monocyte, 6=Monocyte, 7=Other, 8=Other, 9=Endothelial, 10=Fibroblasts, 11=NKT, 12=Other, 13=Pericytes, 14=NKT, 15=Endothelial, 16=Fibroblasts, 17=B-Cell, 18=Monocyte, 19=Endothelial, 20=B-Cell, 21=Fibroblasts, 22=Other, 23=Other, 24=Other, 25=Monocyte, 26=Fibroblasts, 27=Monocyte, 28=Pericytes, 29=Plasma, 30=Pericytes, 31=Endothelial, 32=NKT, 33=Endothelial, 34=Fibroblasts, 35=Monocyte, 36=Monocyte, 37=Other. **D.** Cell type annotations based on canonical marker expression and pericyte gene signature.

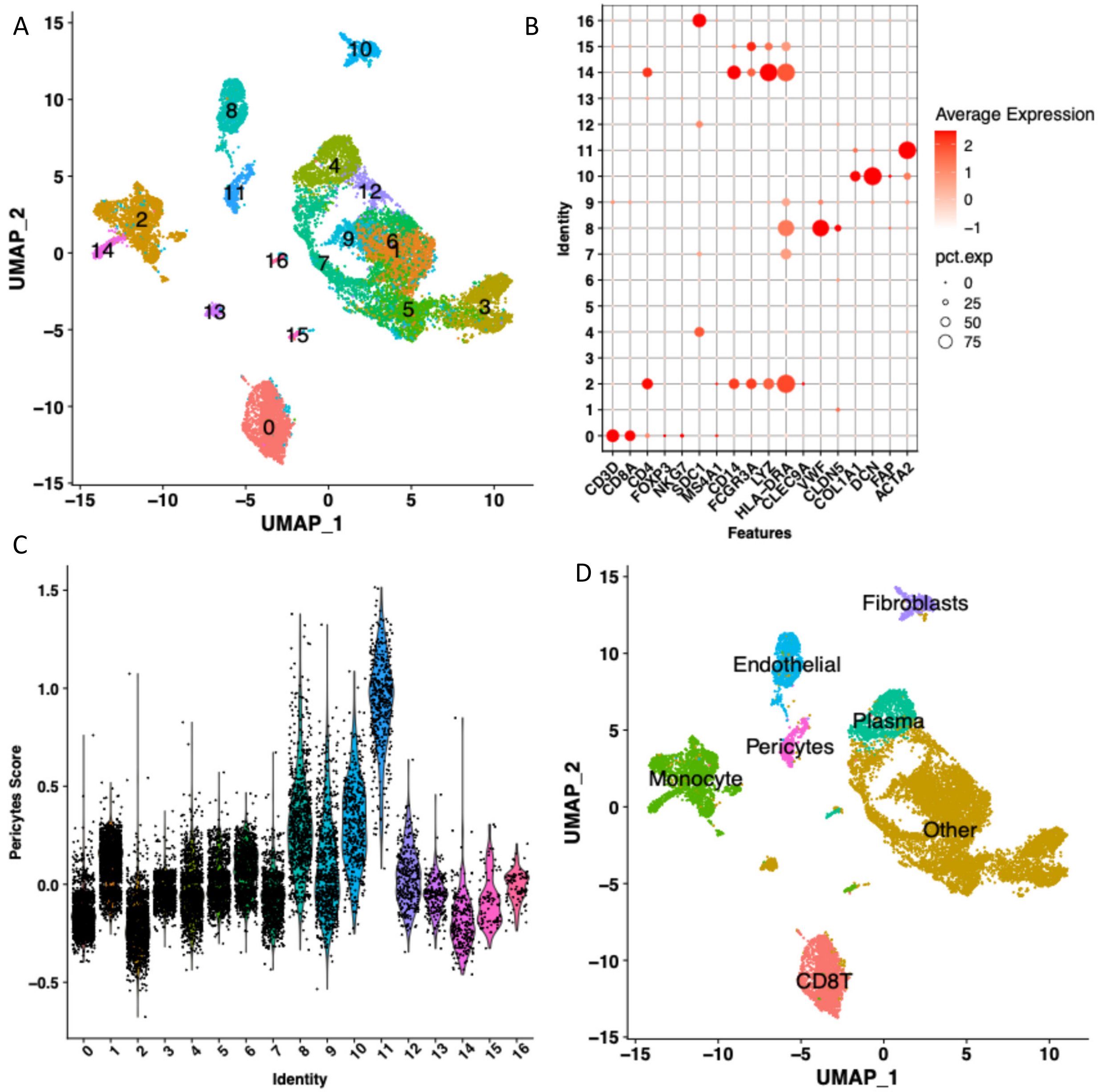

**Supplementary Figure 14. Analysis of prostate cancer dataset (GSE176031): A.** UMAP projection of single-cell transcriptomic data, with cells colored by unsupervised cluster IDs. **B.** Dot plot showing expression of canonical cell type markers across clusters. **C.** Expression of pericyte-related markers across clusters based on module score calculation using genes from Table 1 and the Seurat Package. Cluster ID #11 has high pericytes signature score therefore annotated as pericytes. 0=CD8T, 1=Other, 2=Monocyte, 3=Other, 4=Plasma, 5=Other, 6=Other, 7=Other, 8=Endothelial, 9=Other, 10=Fibroblasts, 11=Pericytes, 12=Other, 13=Other, 14=Monocyte, 15=Monocyte, 16=Plasma. **D.** Cell type annotations based on canonical marker expression and pericyte gene signature.

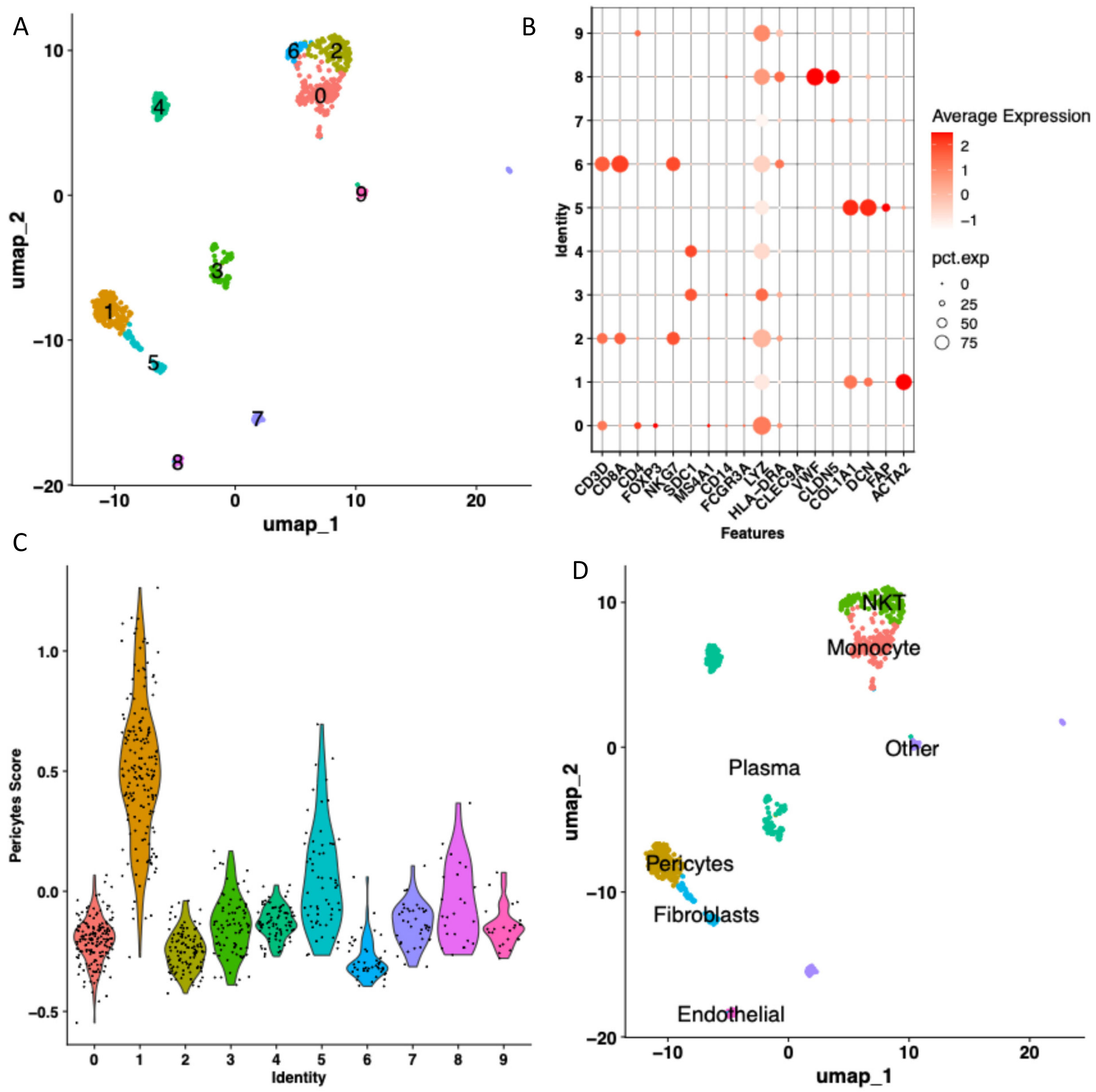

**Supplementary Figure 15. Analysis of head & neck cancer dataset (GSE103322): A.** UMAP projection of single-cell transcriptomic data, with cells colored by unsupervised cluster IDs. **B.** Dot plot showing expression of canonical cell type markers across clusters. **C.** Expression of pericyte-related markers across clusters based on module score calculation using genes from Table 1 and the Seurat Package. Cluster ID #1 has high pericytes signature score therefore annotated as pericytes. 0=Monocyte, 1=Pericytes, 2=NKT, 3=Plasma, 4=Plasma, 5=Fibroblasts, 6=NKT, 7=Other, 8=Endothelial, 9=Other. **D.** Cell type annotations based on canonical marker expression and pericyte gene signature.

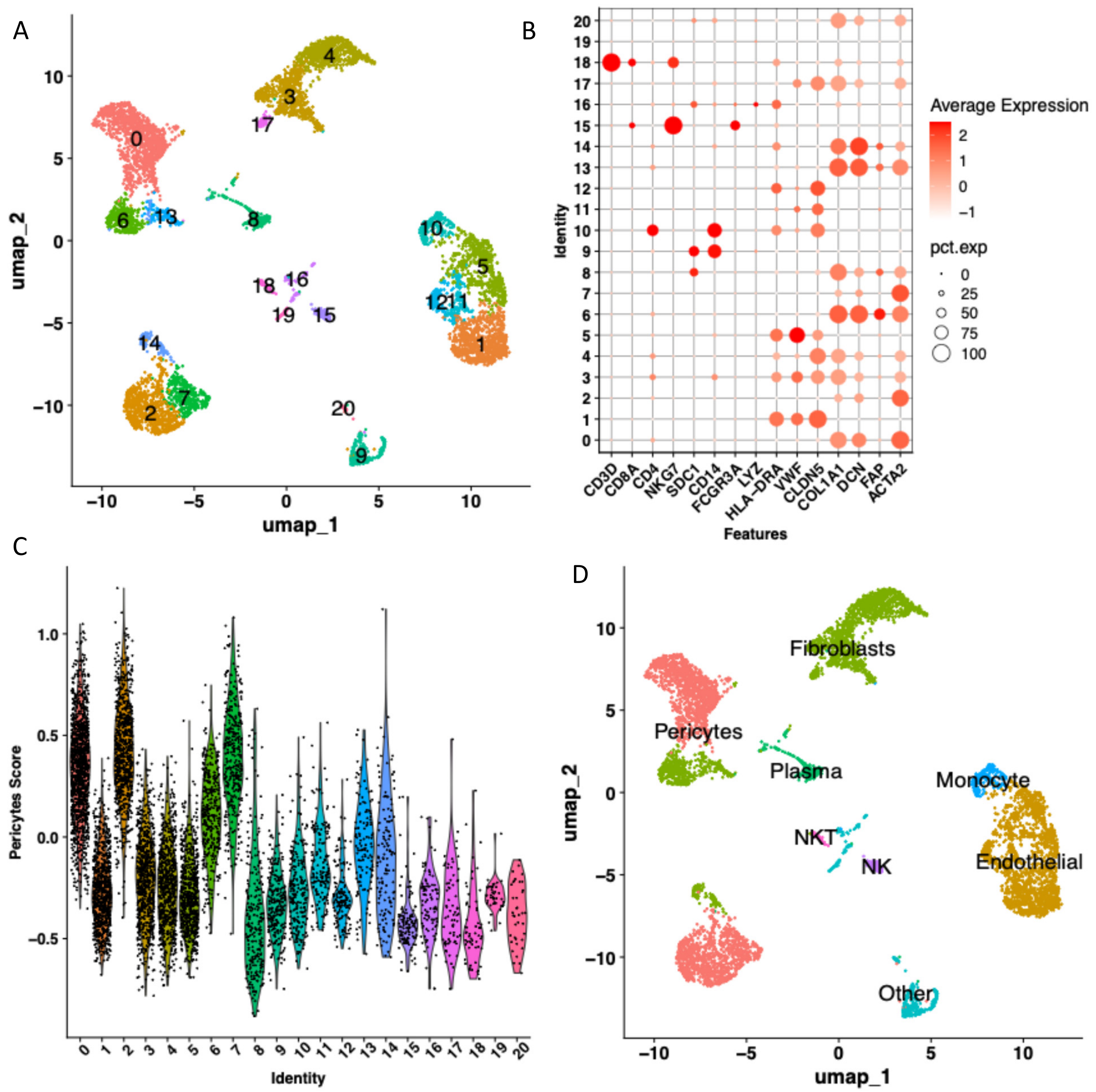

**Supplementary Figure 16. Analysis of cholangiocarcinoma dataset (GSE142784): A.** UMAP projection of single-cell transcriptomic data, with cells colored by unsupervised cluster IDs. **B.** Dot plot showing expression of canonical cell type markers across clusters. **C.** Expression of pericyte-related markers across clusters based on module score calculation using genes from Table 1 and the Seurat Package. Cluster IDs #0, #2 have high pericytes signature score therefore annotated as pericytes. 0=Pericytes, 1=Endothelial, 2=Pericytes, 3=Fibroblasts, 4=Fibroblasts, 5=Endothelial, 6=Fibroblasts, 7=Pericytes, 8=Plasma, 9=Other, 10=Monocyte, 11=Endothelial, 12=Endothelial, 13=Fibroblasts, 14=Fibroblasts, 15=NK, 16=Other, 17=Fibroblasts, 18=NKT, 19=Other, 20=Other. **D.** Cell type annotations based on canonical marker expression and pericyte gene signature.

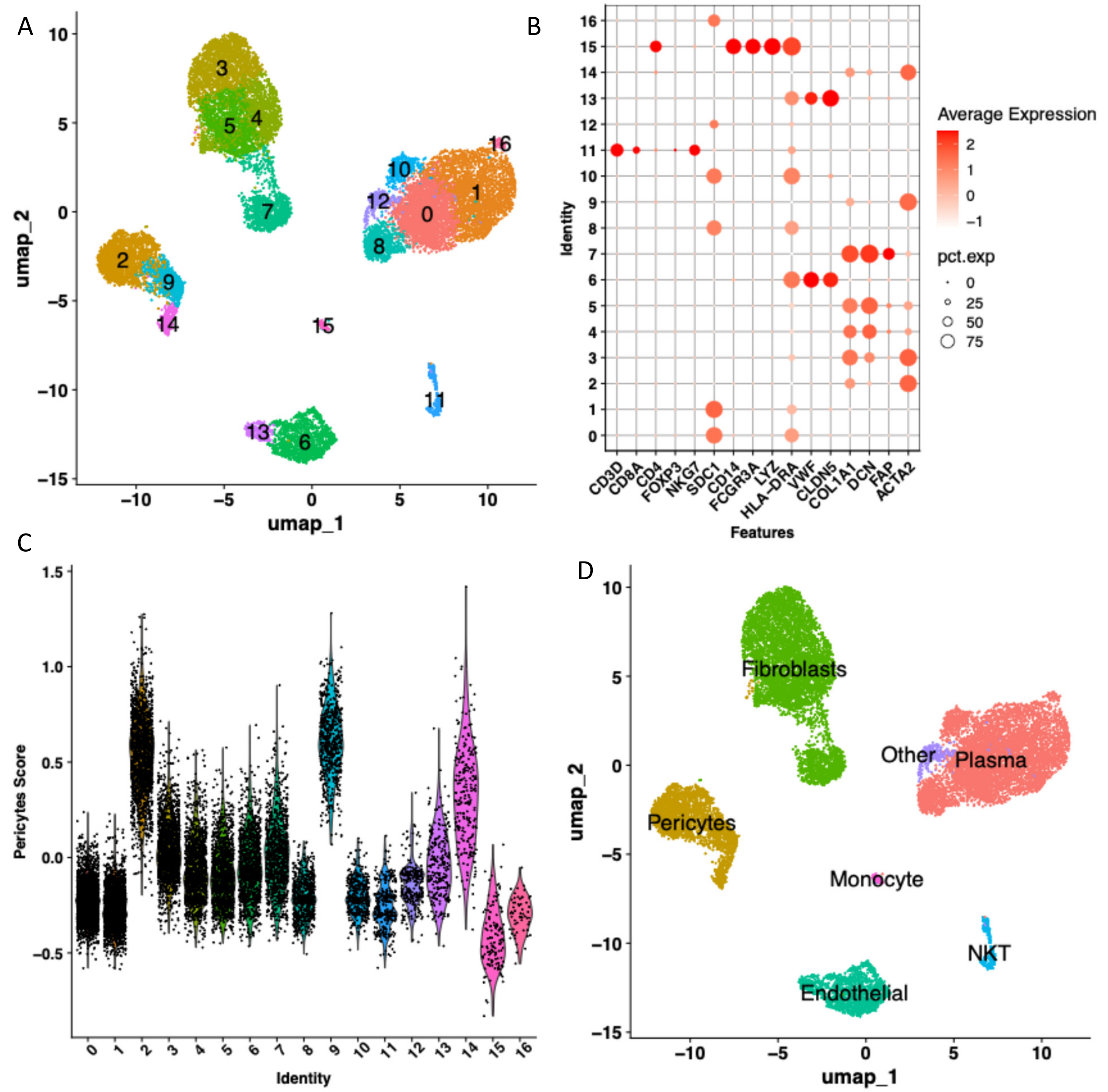

**Supplementary Figure 17. Analysis of cervical cancer dataset (GSE168652): A.** UMAP projection of single-cell transcriptomic data, with cells colored by unsupervised cluster IDs. **B.** Dot plot showing expression of canonical cell type markers across clusters. **C.** Expression of pericyte-related markers across clusters based on module score calculation using genes from Table 1 and the Seurat Package. Cluster IDs #2, #9, #14 have high pericytes signature score therefore annotated as pericytes. 0=Plasma, 1=Plasma, 2=Pericytes, 3=Fibroblasts, 4=Fibroblasts, 5=Fibroblasts, 6=Endothelial, 7=Fibroblasts, 8=Plasma, 9=Pericytes, 10=Plasma, 11=NKT, 12=Other, 13=Endothelial, 14=Pericytes, 15=Monocyte, 16=Plasma. **D.** Cell type annotations based on canonical marker expression and pericyte gene signature.

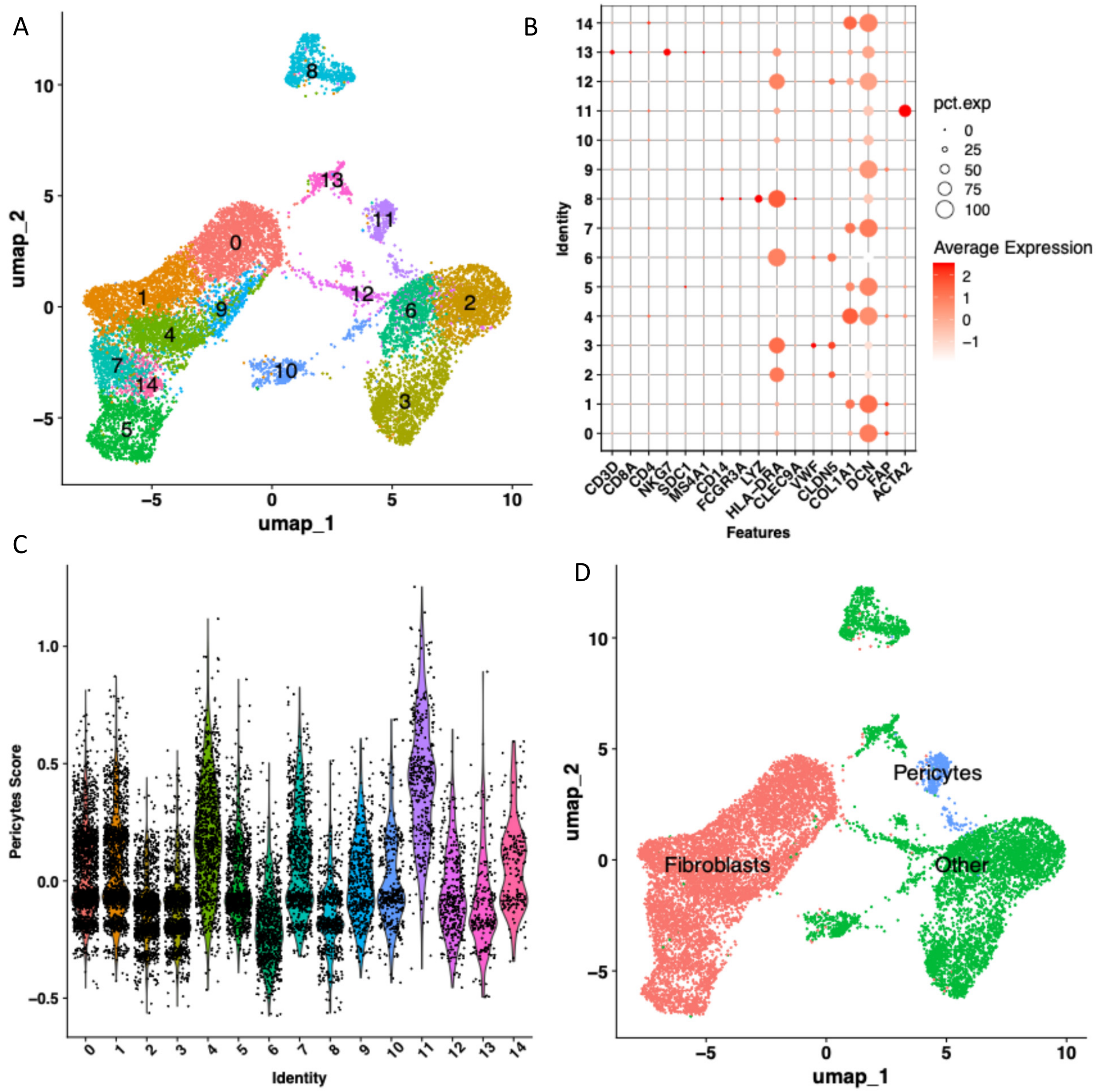

**Supplementary Figure 18. Analysis of neurofibroma dataset (GSE163028): A.** UMAP projection of single-cell transcriptomic data, with cells colored by unsupervised cluster IDs. **B.** Dot plot showing expression of canonical cell type markers across clusters. **C.** Expression of pericyte-related markers across clusters based on module score calculation using genes from Table 1 and the Seurat Package. Cluster ID #11 has high pericytes signature score therefore annotated as pericytes. 0=Fibroblasts, 1=Fibroblasts, 2=Other, 3=Other, 4=Fibroblasts, 5=Fibroblasts, 6=Other, 7=Fibroblasts, 8=Other, 9=Fibroblasts, 10=Other, 11=Pericytes, 12=Other, 13=Other, 14=Fibroblasts. **D.** Cell type annotations based on canonical marker expression and pericyte gene signature.

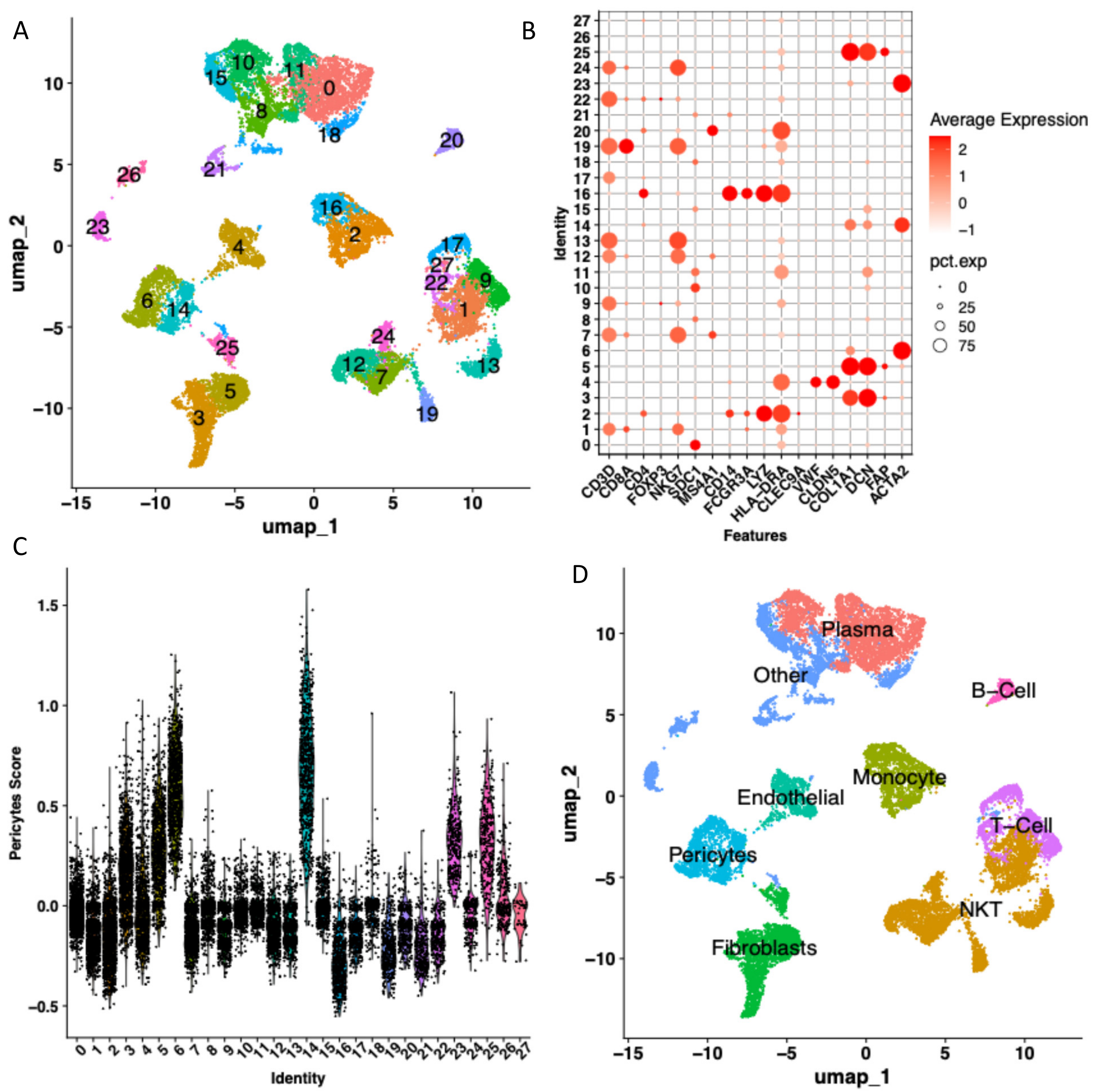

**Supplementary Figure 19. Analysis of lymphoma dataset (GSE128531): A.** UMAP projection of single-cell transcriptomic data, with cells colored by unsupervised cluster IDs. **B.** Dot plot showing expression of canonical cell type markers across clusters. **C.** Expression of pericyte-related markers across clusters based on module score calculation using genes from Table 1 and the Seurat Package. Cluster IDs #6, #14 have high pericytes signature score therefore annotated as pericytes. 0=Plasma, 1=NKT, 2=Monocyte, 3=Fibroblasts, 4=Endothelial, 5=Fibroblasts, 6=Pericytes, 7=NKT, 8=Other, 9=T-Cell, 10=Plasma, 11=Plasma, 12=NKT, 13=NKT, 14=Pericytes, 15=Other, 16=Monocyte, 17=T-Cell, 18=Other, 19=NKT, 20=B-Cell, 21=Other, 22=T-Cell, 23=Other, 24=NKT, 25=Fibroblasts, 26=Other, 27=Other. **D.** Cell type annotations based on canonical marker expression and pericyte gene signature.

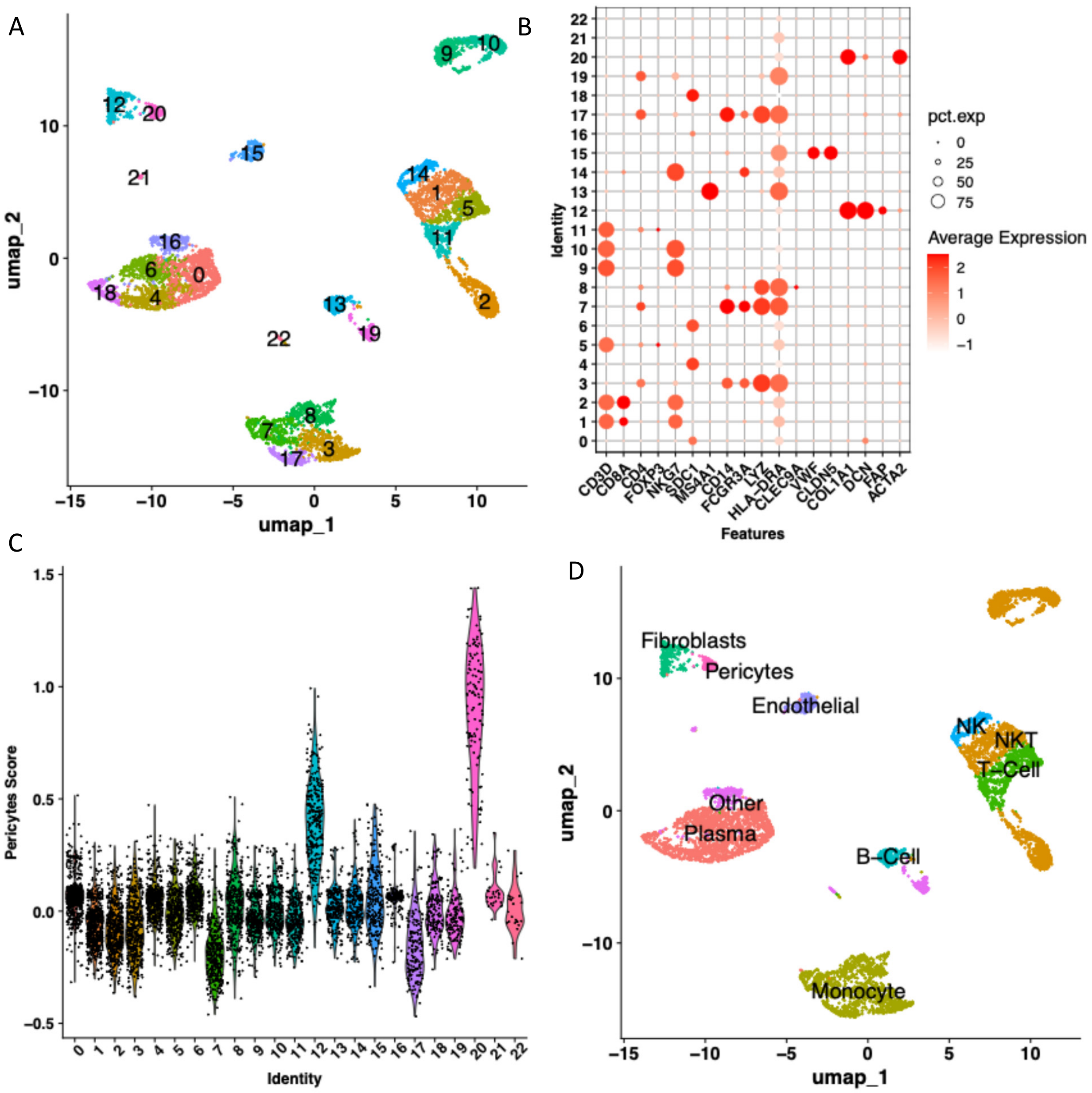

**Supplementary Figure 20. Analysis of lymphoma dataset (GSE147944): A.** UMAP projection of single-cell transcriptomic data, with cells colored by unsupervised cluster IDs. **B.** Dot plot showing expression of canonical cell type markers across clusters. **C.** Expression of pericyte-related markers across clusters based on module score calculation using genes from Table 1 and the Seurat Package. Cluster ID #20 has high pericytes signature score therefore annotated as pericytes. 0=Plasma, 1=NKT, 2=NKT, 3=Monocyte, 4=Plasma, 5=T-Cell, 6=Plasma, 7=Monocyte, 8=Monocyte, 9=NKT, 10=NKT, 11=T-Cell, 12=Fibroblasts, 13=B-Cell, 14=NK, 15=Endothelial, 16=Other, 17=Monocyte, 18=Plasma, 19=Other, 20=Pericytes, 21=Other, 22=Other. **D.** Cell type annotations based on canonical marker expression and pericyte gene signature.

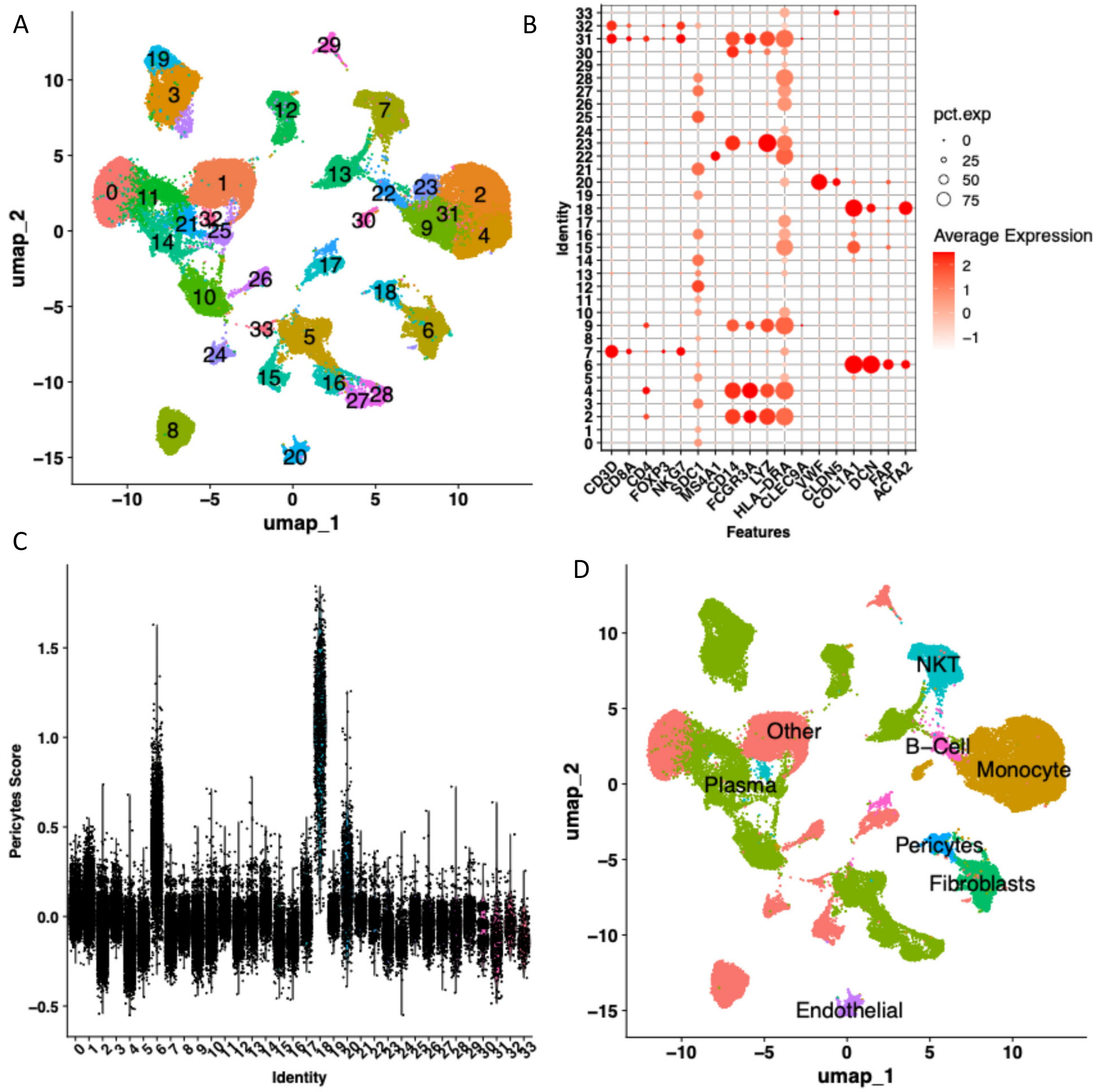

**Supplementary Figure 21. Analysis of lung cancer dataset (GSE148071): A.** UMAP projection of single-cell transcriptomic data, with cells colored by unsupervised cluster IDs. **B.** Dot plot showing expression of canonical cell type markers across clusters. **C.** Expression of pericyte-related markers across clusters based on module score calculation using genes from Table 1 and the Seurat Package. Cluster ID #18 has high pericytes signature score therefore annotated as pericytes. 0=Other, 1=Other, 2=Monocyte, 3=Plasma, 4=Monocyte, 5=Plasma, 6=Fibroblasts, 7=NKT, 8=Other, 9=Monocyte, 10=Plasma, 11=Plasma, 12=Plasma, 13=Plasma, 14=Plasma, 15=Other, 16=Plasma, 17=Other, 18=Pericytes, 19=Plasma, 20=Endothelial, 21=Plasma, 22=B-Cell, 23=Monocyte, 24=Other, 25=Plasma, 26=Other, 27=Plasma, 28=Plasma, 29=Other, 30=Monocyte, 31=Monocyte, 32=NKT, 33=Other. **D.** Cell type annotations based on canonical marker expression and pericyte gene signature.

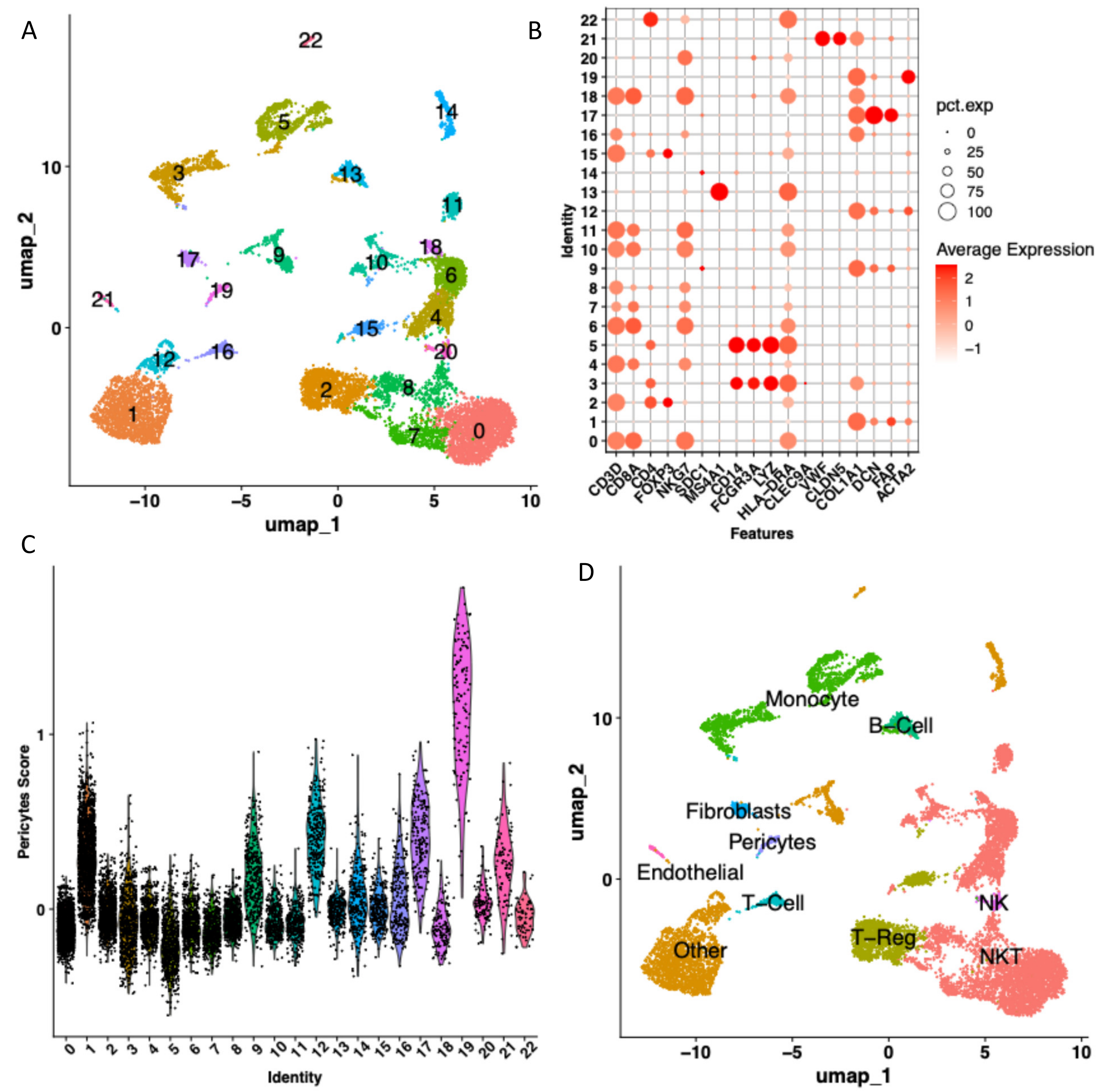

**Supplementary Figure 22. Analysis of thyroid cancer dataset (GSE148673): A.** UMAP projection of single-cell transcriptomic data, with cells colored by unsupervised cluster IDs. **B.** Dot plot showing expression of canonical cell type markers across clusters. **C.** Expression of pericyte-related markers across clusters based on module score calculation using genes from Table 1 and the Seurat Package. Cluster ID #19 has high pericytes signature score therefore annotated as pericytes. 0=NKT, 1=Other, 2=T-Reg, 3=Monocyte, 4=NKT, 5=Monocyte, 6=NKT, 7=NKT, 8=NKT, 9=Other, 10=NKT, 11=NKT, 12=Other, 13=B-Cell, 14=Other, 15=T-Reg, 16=T-Cell, 17=Fibroblasts, 18=NKT, 19=Pericytes, 20=NK , 21=Endothelial, 22=Other. **D.** Cell type annotations based on canonical marker expression and pericyte gene signature.

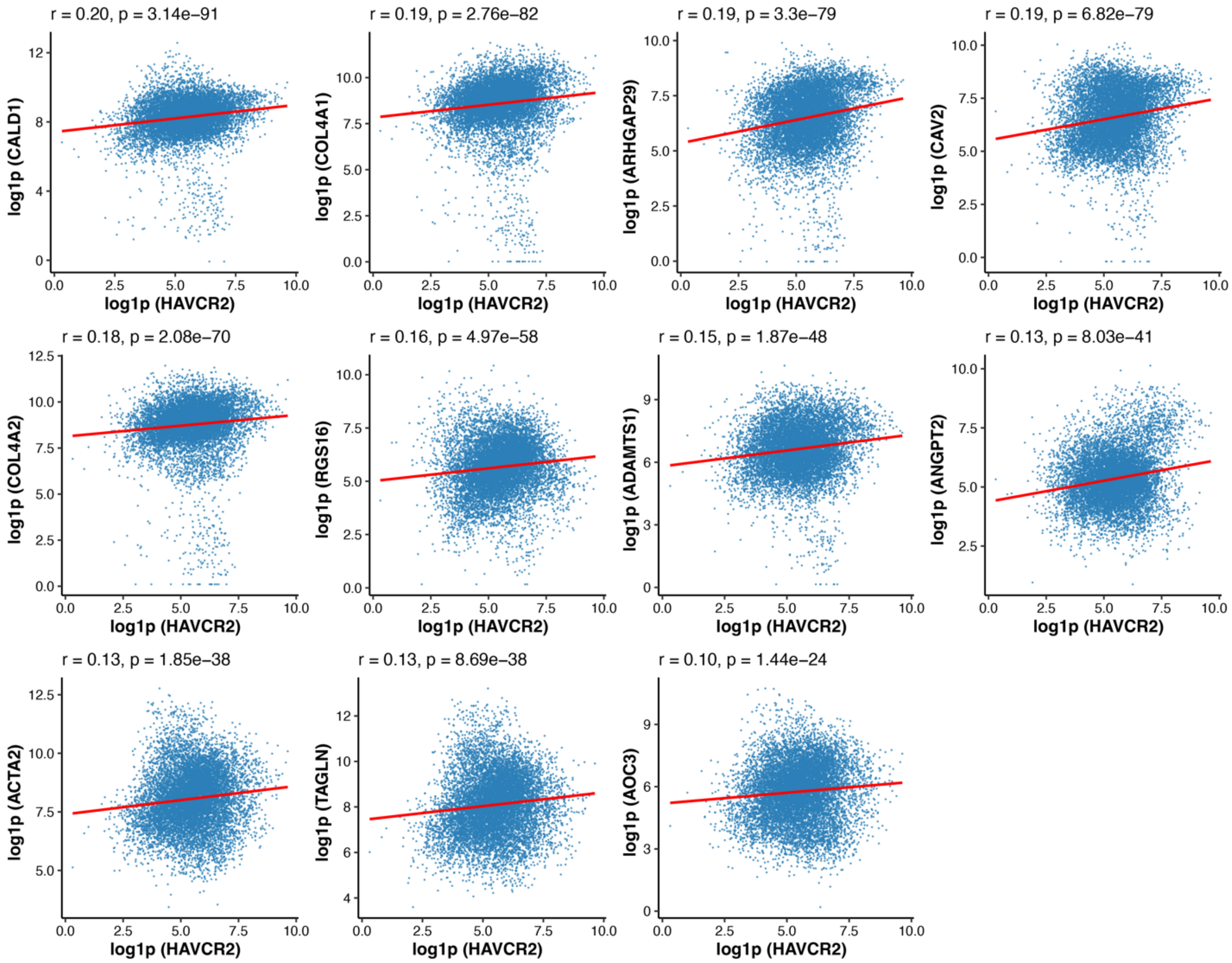

**Supplementary Figure 23.** Scatterplots showing the correlation between Pan-cancer Pericyte Signature genes and the T cell exhaustion marker HAVCR2 (also known as TIM-3). Spearman rank correlation was used to assess the association between gene expression levels.
