## Supplementary material for "Single-cell analysis reveals a universal pericyte signature associated with poor clinical outcome and immune T cell dysfunction in thyroid cancer and other cancers": Suppl. Table 3

**Suppl. Table 3:** Single-cell RNA-sequencing datasets used for the characterization of pericytes across different cancers and benign tumor (neurofibroma).

| Basal Cell Carcinoma GSE123813 | Bladder Urothelial Canceroma GSE130001 | Breast invasive carcinoma-EMTAB8107 |
| --- | --- | --- |
| Cholangiocarcinoma GSE142784 | Colorectal Cancer GSE166555 | Head and neck squamous cell carcinoma GSE103322 |
| Neurofibroma GSE163028 | Non-Hodgkin Lymphoma GSE128531 | Non-Hodgkin Lymphoma GSE147944 |
| Non-small cell lung cancer GSE148071 | Ovarian Serous Cystadenocarcinoma EMTAB8107 | Ovarian Serous Cystadenocarcinoma GSE118828 |
| Ovarian Serous Cystadenocarcinoma GSE130000 | Ovarian Serous Cystadenocarcinoma-GSE154600 | Cervical Squamous Cell Carcinoma and Endocervical Adenocarcinoma GSE168652 |
| Thyroid Cancer  GSE148673 and GSE193581 | Pancreatic adenocarcinoma-CRA001160 | Pancreatic adenocarcinoma GSE154778 |
| Prostate adenocarcinoma GSE137829 | Prostate adenocarcinoma GSE172301 | Prostate adenocarcinoma GSE176031 |
| Skin Cutaneous melanoma GSE134388 |  |  |
